# Inhibition of Soluble Epoxide Hydrolase Ameliorates Cerebral Blood Flow Autoregulation and Cognition in Alzheimer’s Disease and Diabetes-Related Dementia Rat Models

**DOI:** 10.1101/2024.08.30.610540

**Authors:** Chengyun Tang, Jane J. Border, Huawei Zhang, Andrew Gregory, Shan Bai, Xing Fang, Yedan Liu, Shaoxun Wang, Sung Hee Hwang, Wenjun Gao, Gilbert C. Morgan, Jhania Smith, David Bunn, Cameron Cantwell, Karen M. Wagner, Christophe Morisseau, Jun Yang, Seung Min Shin, Philip O’Herron, Zsolt Bagi, Jessica A. Filosa, Yanbin Dong, Hongwei Yu, Bruce D. Hammock, Richard J. Roman, Fan Fan

## Abstract

Alzheimer’s Disease and Alzheimer’s Disease-related dementias (AD/ADRD) pose major global healthcare challenges, with diabetes mellitus (DM) being a key risk factor. Both AD and DM-related ADRD are characterized by reduced cerebral blood flow, although the exact mechanisms remain unclear. We previously identified compromised cerebral hemodynamics as early signs in TgF344-AD and type 2 DM-ADRD (T2DN) rat models. Genome-wide studies have linked AD/ADRD to SNPs in soluble epoxide hydrolase (sEH). This study explored the effects of sEH inhibition with TPPU on cerebral vascular function and cognition in AD and DM-ADRD models. Chronic TPPU treatment improved cognition in both AD and DM-ADRD rats without affecting body weight. In DM-ADRD rats, TPPU reduced plasma glucose and HbA1C levels. Transcriptomic analysis of primary cerebral vascular smooth muscle cells from AD rats treated with TPPU revealed enhanced pathways related to cell contraction, alongside decreased oxidative stress and inflammation. Both AD and DM-ADRD rats exhibited impaired myogenic responses and autoregulation in the cerebral circulation, which were normalized with chronic sEH inhibition. Additionally, TPPU improved acetylcholine-induced vasodilation in the middle cerebral arteries (MCA) of DM-ADRD rats. Acute TPPU administration unexpectedly caused vasoconstriction in the MCA of DM-ADRD rats at lower doses. In contrast, higher doses or longer durations were required to induce effective vasodilation at physiological perfusion pressure in both control and ADRD rats. Additionally, TPPU decreased reactive oxygen species production in cerebral vessels of AD and DM-ADRD rats. These findings provide novel evidence that chronic sEH inhibition can reverse cerebrovascular dysfunction and cognitive impairments in AD/ADRD, offering a promising avenue for therapeutic development.

## INTRODUCTION

Alzheimer’s Disease and Alzheimer’s Disease-Related Dementias (AD/ADRD) affect 6.9 million Americans at age over 65 years in 2024, with projections indicating a potential rise to 13 million by 2050.^1^ This staggering increase underscores the urgent need for effective pharmacotherapies for AD/ADRD. However, despite extensive efforts, such treatments remain elusive, emphasizing the critical importance of gaining a deeper understanding of the underlying cellular and molecular mechanisms driving the onset and progression of the diseases.^2,3^ Brain hypoperfusion has been recognized as one of the primary contributors to AD/ADRD-related neuronal damage, occurring prior to the onset of amyloid beta (Aβ) and tau pathologies and preceding cognitive decline in both AD and ADRD, which is often associated with hypertension and diabetes (DM).^4–8^ Currently, none of the U.S. Food and Drug Administration (FDA)-approved treatments have targeted cerebral vasculature, and AD/ADRD has not been proven curable largely due to an incomplete understanding of the precise underlying mechanisms.^2,3^

Reduced brain perfusion in AD/ADRD has been reported to arise from impaired cerebral blood flow (CBF) autoregulation, blood-brain barrier (BBB) damage, neurovascular uncoupling, and diminished venous and neurovascular-glymphatic function.^6,9–13^ Inflammation and oxidative stress are intricately linked to cerebral vascular dysfunction in AD/ADRD, involving diverse mechanisms and phenotypes.^14,15^ The myogenic response is pivotal for maintaining CBF autoregulation. Mural cells situated on the vessel wall, including vascular smooth muscle cells (VSMCs) and pericytes expressing alpha-smooth muscle actin (α-SMA), are instrumental in modulating the myogenic response of cerebral arteries and arterioles, including the middle cerebral arteries (MCAs) and penetrating arterioles (PAs).^12,16–18^ In AD/ADRD patients and animal models, the VSMCs shift from their typical contractile state to a phenotype resembling that of macrophages. This shift is accompanied by the presence of additional neurodegenerative markers, highlighting the central role of these VSMCs in driving disease progression.^19,20^ The loss of contractility in VSMCs, in turn, can negatively affect the ability of CBF autoregulation in AD/ADRD.^11,21–23^ We previously observed impaired cerebral hemodynamics preceding cognitive deficits in both TgF344-AD and type 2 DM-related ADRD (T2DN) rat models.^7,24,25^ The diminished myogenic response and CBF autoregulation, as well as compromised functional hyperemia, are associated with reduced contractility in cerebral VSMCs and α-SMA positive pericytes, enhanced production of reactive oxidative species (ROS) and mitochondrial-derived superoxide, and diminished adenosine triphosphate production and elevated inflammation.^7,11,24,25^

The soluble epoxide hydrolase (sEH) is an enzyme that transforms arachidonic acid-derived epoxyeicosatrienoic acids (EETs) to corresponding diols dihydroxyeicosatrienoic acids (DHETs) and linoleic acid-derived epoxyoctadecenoic acids (EpOMEs) to dihydroxyoctadecenoic acids (DiHOMEs).^26^ The sEH enzyme is widely distributed in neurons, glial cells, VSMCs, and endothelial cells in the brain.^26–28^ Recent studies suggest that sEH inhibitors show promise in treating a range of conditions in animal models, including cardiovascular diseases,^29–31^ renal diseases,^31,32^ neurodegenerative diseases,^33,34^ and pain-related disorders. ^35–37^ Cognitive function was also improved in cerebral hypoperfusion animal models treated with sEH inhibitors, though the underlying vascular mechanisms remain not fully understood.^38,39^ In the present study, we compared the transcriptomic profiling of primary VSMCs isolated from AD rats treated with a highly selective sEH inhibitor 1-(1-Propanoylpiperidin-4-yl)-3-[4-(trifluoromethoxy)phenyl]urea (TPPU) and evaluated the impact of sEH inhibition on myogenic response, CBF autoregulation, VSMC contractility, oxidative stress, and cognition in AD and DM-related ADRD rats.

## MATERIALS AND METHODS

### General

Experiments were conducted in age-matched diabetic T2DN (DM or DM-ADRD) vs. Sprague Dawley (SD) and TgF344-AD (AD) vs. Fischer 344 (F344) rats. TPPU was kindly provided by Dr. Hammock. TPPU was dissolved in 100% polyethylene glycol, molecular weight 400 (PEG-400; 91893, MilliporeSigma, Burlington, MA) then further diluted in drinking water to achieve a final PEG-400 concentration of 1%. The concentration of TPPU in the PEG-400 solution was adjusted based on body weight and daily water intake, with the values averaged over a pre-measured three-day period. The rats were treated with either a vehicle (1% PEG-400 in drinking water) or TPPU (1 mg/kg/day in the vehicle) ^40–42^ for 9 weeks, beginning at 15 months of age for SD and T2DN rats, and 9 months of age for F344 and AD rats. The rats were sourced from inbred colonies at the University of Mississippi Medical Center and Augusta University. Standard laboratory animal conditions were maintained, including a 12-hour light-dark cycle and *ad libitum* access to food and water throughout the study. The animal care facilities hold accreditation from the American Association for the Accreditation of Laboratory Animal Care. All animal experiments adhered to protocols approved by the Institutional Animal Care and Use Committees. Body Weight, Plasma Glucose, and Glycated Hemoglobin A1c (HbA1c) Measurements

Conscious rats were weighed every three weeks, and blood samples were collected from the tail vein three hours after the lights-on cycle. Following our previously described protocols, ^43^ plasma glucose levels were determined using a Contour Next glucometer (Ascensia Diabetes Care, Parsippany, NJ), and HbA1c levels were assessed with a PTS Diagnostics A1CNow+ device (Fisher Scientific, Hampton, NH).

### Eight-Arm Water Maze

Spatial learning and short- and long-term memory were assessed with an eight-arm water maze.^17^ Rats were initially trained to locate a platform marked with a visual cue in one of the eight arms of a pool for escape. Subsequently, four memory trials were conducted at 2- and 24-hour post-training, each consisting of four consecutive swims. The latency to reach the platform was recorded.^7,25^

### Bulk RNA-seq Analysis

Primary VSMCs were obtained from the MCAs of 3-week-old male AD and F344 rats. The isolation process involved digestion with protease dithiothreitol (2 mg/mL) and papain (22.5 U/mL), followed by collagenase (250 U/mL), elastase (2.4 U/mL), and trypsin inhibitor (10,000 U/mL), as previously described in our report. ^17,22,43,44^ Early passages (P2-4) of cells from 3 rats were cultured in Dulbecco’s modified Eagle’s medium (DMEM, Thermo Fisher Scientific, Waltham, MA) supplemented with 10% fetal bovine serum (Thermo Scientific) at 37°C with 5% CO_2_. The cells were treated with TPPU (10 µM) or vehicle (0.1% DMSO) for 40 hours. The dosage and duration of TPPU treatments in VSMCs were established through preliminary cell contraction studies. ^45^ Total RNA from vehicle- or TPPU-treated AD cerebral VSMCs was extracted using Trizol reagent (Thermo Fisher Scientific) followed with Purelink RNA mini kit (Thermo Fisher Scientific). RNA quality assessment, RT-qPCR, library preparation, and RNA sequencing were all performed in the Integrated Genomics Shared Resource at the Georgia Cancer Center. High-throughput RNA sequencing raw reads were obtained. The initial processing of the raw data involved trimming and removing adapters using Cutadapt 4.0.^46^ Sequences were aligned to the rat genome (rn7) using STAR 2.7.10a, ^47^ and read counts were obtained. The differential expression and enrichment analysis were performed using DESeq2^48^ and clusterProfiler, ^49^ respectively. The KEGG (Kyoto Encyclopedia of Genes and Genomes) database was used for enrichment analysis.

### Cerebral Vascular Myogenic Reactivities

#### Vessel Preparation

The MCAs and PAs were isolated following our established protocol.^23,50,51^ Rats were euthanized under 4% isoflurane anesthesia. Brains were rapidly removed and placed in a dish containing ice-cold calcium-free physiological salt solution (PSS_0Ca_), composing 119 mM NaCl, 4.7 mM KCl, 1.17 mM MgSO_4_, 0.03 mM EDTA, 18 mM NaHCO_3_, 5 mM HEPES, 1.18 mM NaH_2_PO_4_, and 10 mM glucose (pH 7.4).^52–54^ The branch-free M2 segments of the MCAs and the downstream PAs were dissected in ice-cold PSS_0Ca_ supplemented with 1% bovine serum albumin.

#### Impact of Chronic sEH Inhibition on Myogenic Response in Cerebral Vasculature

The branch-free segment of the MCA-M2 was cannulated onto glass micropipettes (1B120-6, World Precision Instruments, Sarasota, FL) and mounted in a pressure myograph chamber (Living System Instrumentation, Burlington, VT).^43,55^ The chamber was filled with PSS_Ca_ and aerated with a gas mixture of 21% O_2_, 5% CO_2_, and 74% N_2_ to maintain pH 7.4 at 37 °C. The pressure myograph chamber was connected to an IMT-2 inverted microscope (Olympus, Center Valley, PA) outfitted with a digital camera (MU1000, AmScope, Irvine, CA). The cannulated vessels were adjusted to their *in situ* length at an intraluminal pressure of 40 mmHg and allowed to equilibrate for 30 minutes to develop spontaneous tone. ^51,56,57^ Vascular viability was confirmed by ensuring a vasoconstrictive response of more than 15% to 60 mM KCl in PSS containing 1.6 mM CaCl_2_ (PSS_Ca_) within the bath.^58^ Vessels failing to meet this criterion were excluded from the study. After preconditioning, the inner diameters (IDs) of MCAs were measured in rats treated for 9 weeks with either vehicle or TPPU. The myogenic response was assessed by recording the pressure-diameter relationships in response to transmural pressures ranging from 40 to 180 mmHg in increments of 20 mmHg. Similarly, the myogenic response of PAs in rats treated for 9 weeks with either vehicle or TPPU was assessed by recording the pressure-diameter relationships in response to transmural pressures ranging from 10 to 60 mmHg in increments of 10 mmHg.

#### Impact of Chronic sEH Inhibition on Acetylcholine (Ach)-Induced Vasodilation

To assess endothelial-related vasodilation, the MCAs were pressurized to 40 mmHg and precontracted with 5-HT (10^-4^ M). The changes of IDs of the MCA in vehicle or TPPU-treated DM-ADRD rats in response to Ach (10^-8^ to 10^-4^ M) were evaluated.

#### Impact of Acute sEH Inhibition on Myogenic Response in Cerebral Vasculature

These experiments were conducted using male rats aged 24-36 weeks for TgF344- AD vs. F344 and 24-48 weeks for T2DN vs. SD, chosen based on our previous findings indicating the development of cerebral vascular and cognitive impairments in these AD and DM-ADRD models at these time points.^7,25^ Freshly isolated intact MCAs were pressurized from 40 to 120 mmHg. The vessels were superfused with either vehicle (0.1% DMSO) or TPPU (1 µM and 10 µM) in the bath at 120 mmHg. The TPPU dosage was established via a pilot study, which tested 0.1, 1, and 10 µM TPPU in DMSO in SD rats. Data for the 0.1 µM concentration are not presented. ID changes were recorded every 5 minutes for 30 minutes.

### CBF Autoregulation

The impact of chronic inhibition of sEH on CBF autoregulation was accessed using laser Doppler flowmetry with our established protocol.^17,25^ Rats underwent tracheal, femoral vein, and artery cannulation after anesthesia induction with inactin (50 mg/kg; *i.p.*) and ketamine (30 mg/kg; *i.m.*). The head was immobilized in a stereotaxic device (Stoelting, Wood Dale, IL), and an end-tidal P_CO2_ of 35 mmHg was maintained using a CO_2_ Analyzer (CAPSTAR-100, CWE Inc., Ardmore, PA). Mean arterial pressure (MAP) was measured using a pressure transducer connected to the cannulated femoral artery. A thin, translucent close cranial window was created 2 mm posterior and 6 mm lateral to the bregma using a low-speed air drill. A fiber-optic probe (91-00124, Perimed Inc., Las Vegas, NV) connected to a laser Doppler flowmetry device (PF5010, Perimed Inc.) was positioned over the cranial window. Baseline CBF was recorded at 100 mmHg. Perfusion pressure was then increased to 180 mmHg in increments of 20 mmHg by infusing phenylephrine (P6126; MilliporeSigma) at doses of 0.5 to 5 µg/min. After recording the response to pressure elevations, a new baseline CBF at 100 mmHg was established by withdrawing phenylephrine. CBF was recorded at each MAP level down to 40 mmHg through graded hemorrhage in 20 mmHg steps.

### VSMC Contraction Assay

The myogenic response, an intrinsic property of VSMCs, plays a crucial role in regulating CBF autoregulation. The effect of sEH inhibition was evaluated on the contractile capabilities of cerebral VSMCs in F344 and AD rats.

Primary cerebral VSMCs were isolated from the MCA of F344 and AD rats using our established protocol. ^18,22^ The rats were euthanized with isoflurane, and the dissected MCAs were incubated with a combination of protease papain (22.5 U/mL) and dithiothreitol (2 mg/mL), followed by elastase (2.4 U/mL), collagenase (250 U/mL), and trypsin inhibitor (10,000 U/mL). TPPU was initially dissolved in 100% DMSO to a concentration of 100 µM (1,000 X). The contractile capability of cerebral VSMCs from AD rats, treated with either vehicle (0.1% DMSO) or TPPU (0.1 µM), was assessed using a collagen gel-based assay kit (CBA-201, Cell Biolabs, San Diego, CA), as previously described.^21,22^

Cells (2 x 10^6^ cells/mL), suspended in Dulbecco’s Modified Eagle’s Medium (Thermo Scientific, Waltham, MA) supplemented with 20% fetal bovine serum and pre-treated with vehicle (0.1% DMSO) or 0.1 µM TPPU for 30 minutes, were mixed with four times the volume of a collagen gel working solution on ice. The cell-gel mixture (500 μL) in each well of a 24-well plate was initially incubated at 37°C for 1 hour. Subsequently, 1 mL of additional culture medium containing either vehicle (0.1% DMSO) or TPPU (0.1 µM) was added and incubated for 24 hours at 37°C in a 5% CO_2_ atmosphere to induce contractile stress. To initiate contraction, the stressed matrix was dislodged from the well wall using a sterile needle. Gel sizes were measured at this point as control values. Following initial contraction, 1 mL of DMEM containing vehicle (0.1% DMSO) or TPPU (0.1 µM) was applied to the collagen gel. Changes in gel size were then imaged and quantified at 120 minutes.

### Determination of ROS Production

ROS production in MCAs of SD and F344 rats, as well as T2DN and AD rats treated with either vehicle or TPPU, was evaluated using dihydroethidium (DHE, 10 μM) staining and the MitoSOX™ Red Mitochondrial Superoxide Indicator kit (10 µM, M36008, Thermo Scientific), according to our established protocols. ^22,23^ After staining, the vessels were fixed with 3.7% paraformaldehyde, mounted with VECTASHIELD Antifade Mounting Medium (H-1000, Vector Laboratories, Burlingame, CA), and imaged using a Nikon Eclipse 55i fluorescence microscope (Nikon, Melville, NY) with excitation/emission wavelengths of 480/576 nm (DHE) and 405/590 nm (MitoSOX). Quantification was performed by comparing the mean intensities of red fluorescence in vessels from control and drug-treated animals using Image J software.

### Statistics

The results are expressed as mean values ± standard error of the mean (SEM). Differences among continuously measured groups were analyzed using a two-way ANOVA for repeated measures, followed by a Holm-Sidak *post hoc* test. To assess differences between two groups or treatments, paired or unpaired t-tests were used, or a one-way ANOVA followed by a Tukey *post hoc* test was performed. Statistical analyses were conducted using GraphPad Prism 10 (GraphPad Software, Inc., La Jolla, CA). Statistical significance was defined as *p* < 0.05 for most analyses. For the bulk RNA-seq analysis, significance was determined using a threshold of *q* < 0.05, where q-values represent False Discovery Rate (FDR)-adjusted *p* -values.

## RESULTS

### Impact of Chronic sEH Inhibition on Body Weight, Plasma Glucose, and HbA1c Levels in AD/ADRD Rats

As depicted in **Figure 1** (upper panel), similar to our previous findings in 18-month-old male DM rats, ^25,43^ the baseline body weights of 15-month-old male DM rats (408.0 ± 6.5 g, n = 12, *p* < 0.0001) were significantly lower than those of non-diabetic controls (487.0 ± 18.8 g, n = 6). Body weights remained consistently lower in TPPU-treated DM-ADRD rats compared to SD rats, with no further reduction observed. Sustained sEH inhibition for 9 weeks reduced elevated plasma glucose (from 316.4 ± 14.8 mg/dL to 273.3 ± 16.1 mg/dL, n = 12, *p* < 0.0001) and HbA1c levels (from 10.5 ± 0.3% to 9.9 ± 0.4%, n = 12, *p* = 0.0013) in DM rats (**Figure 1**, middle and lower panels). Notably, plasma glucose and HbA1c levels showed further reductions starting from the 3-week time point of TPPU treatment in DM-ADRD rats, continuing through the 6-week and 9-week treatments. In contrast, TPPU treatment did not result in any alteration in body weight, glucose, and HbA1c levels in AD rats throughout the 9-week treatment period.

**Figure 1.**
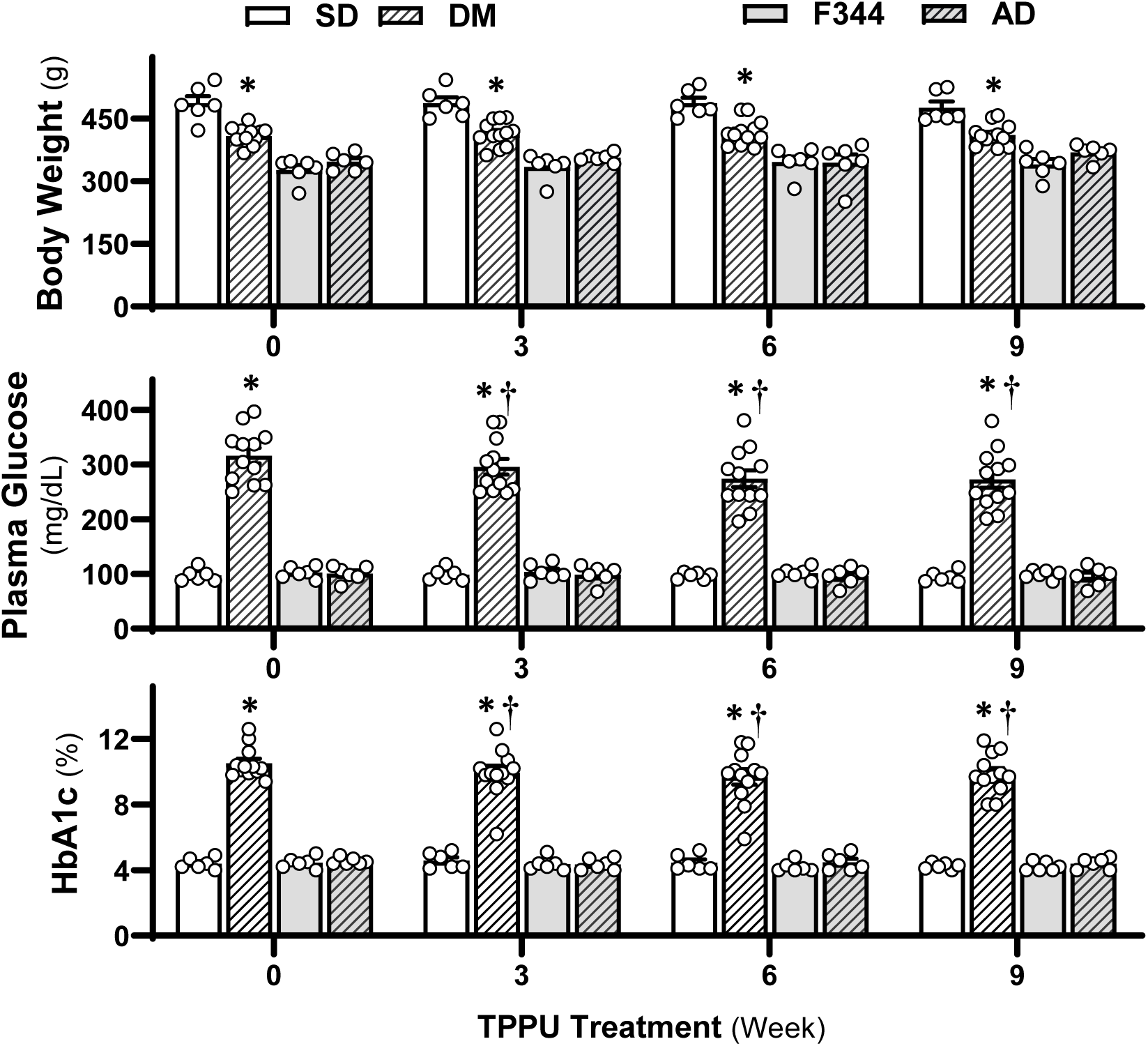
Impact of Chronic sEH Inhibition on Body Weight, Plasma Glucose, and Glycated Hemoglobin A1c (HbA1c) Levels in AD/ADRD Rats. Comparison of body weight (upper panel), plasma glucose (middle panel), and HbA1c (lower panel) in age-matched diabetic T2DN (DM) vs. Sprague Dawley (SD) and TgF344-AD (AD) vs. F344 rats. These rats were treated with either a vehicle (1% PEG-400 in drinking water) or a sEH inhibitor TPPU (1 mg/kg/day) for 9 weeks. Mean values ± SEM are presented, with 6-12 rats per group. * denotes *p* < 0.05 from the corresponding values in AD or DM-ADRD rats compared to their respective control rats. **†** denotes *p* < 0.05 in corresponding values from TPPU-treated rats compared to untreated rats (week 0) within the same strain.

### Inhibition of sEH Improves Cognition in AD/ADRD rats

Spatial navigation abilities and hippocampus-dependent memory cognition were evaluated with an eight-arm water maze study. As presented in **Figure 2**, in vehicle-treated groups, DM-ADRD (upper panel) and AD (lower panel) rats took significantly longer to escape compared with related wild-type controls, indicating cognitive impairments. However, TPPU treatment reversed cognitive decline in both AD and DM-ADRD rats. In contrast, cognitive function was not altered in TPPU-treated controls.

**Figure 2.**
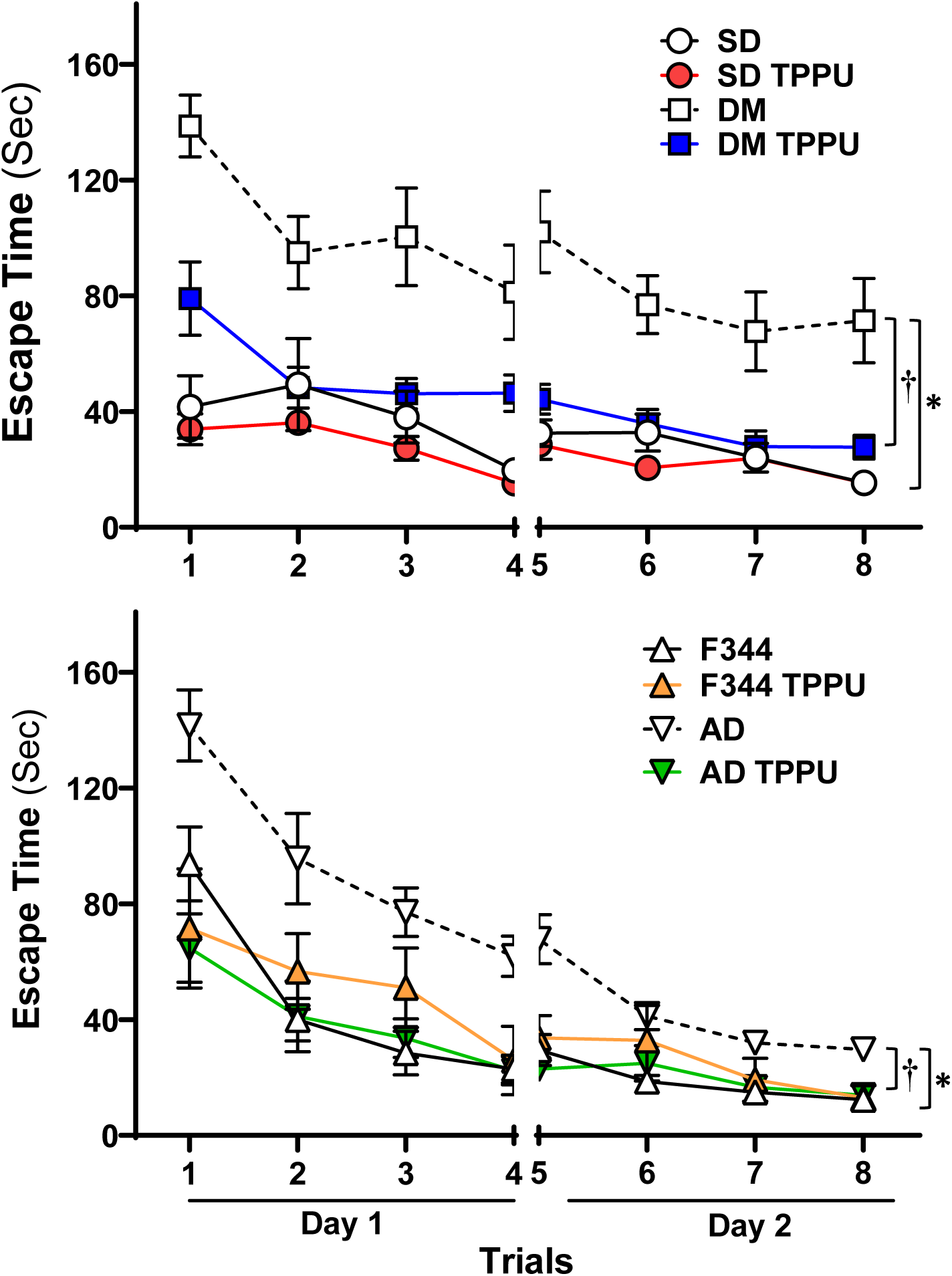
Inhibition of sEH Improves Cognition in AD/ADRD rats. Spatial navigation abilities and hippocampus-dependent memory cognition were evaluated with an eight-arm water maze. Mean values ± SEM are presented, with 6-12 rats per group. * denotes *p* < 0.05 from the corresponding values in AD or DM-ADRD rats compared to their respective control rats. **†** denotes *p* < 0.05 in corresponding values from TPPU-treated rats compared to vehicle-treated rats within the same strain.

### Bulk RNA-seq Analysis

Transcriptomic profiling analysis revealed that 2,178 genes are upregulated (*p* < 0.05), and 2,202 genes are downregulated (*p* < 0.05) in primary cerebral VSMCs from AD rats compared to F344 control rats. In contrast, 3,218 genes are upregulated (*p* < 0.05), and 3,330 genes are downregulated (*p* < 0.05) in AD rats treated with TPPU compared to those receiving the vehicle control (see **volcano plots in Figures 3A, B**). The top pathways significantly altered (FDR-adjusted *p* < 0.05) by KEGG pathway enrichment analysis in primary cerebral VSMCs from AD rats compared to F344 control rats, and in TPPU-treated AD rats compared to vehicle-treated AD rats, are displayed in **bar graphs in Fig. 3A and 3B**. These pathways are essential for regulating cellular contraction (e.g., Vascular Smooth Muscle Contraction, Regulation of Actin Cytoskeleton, AGE-RAGE Signaling Pathway, Focal Adhesion, cAMP Signaling Pathway), oxidative stress (e.g., Oxidative Phosphorylation, AMPK Signaling Pathway, cAMP Signaling Pathway, cGMP-PKG Signaling Pathway, Fluid Shear Stress, Pathways of Neurodegeneration), and inflammation (e.g., AGE-RAGE Signaling Pathway), all of which are crucial mechanisms in the regulation of CBF Relative gene expression levels crucial for regulating these processes are compared in **Fig. 3C and 3D**. Generally, AD cells exhibit reduced expression of genes related to cellular contraction and increased expression of genes associated with oxidative stress and inflammation, while TPPU treatment in AD cells reverses these effects.

**Figure 3.**
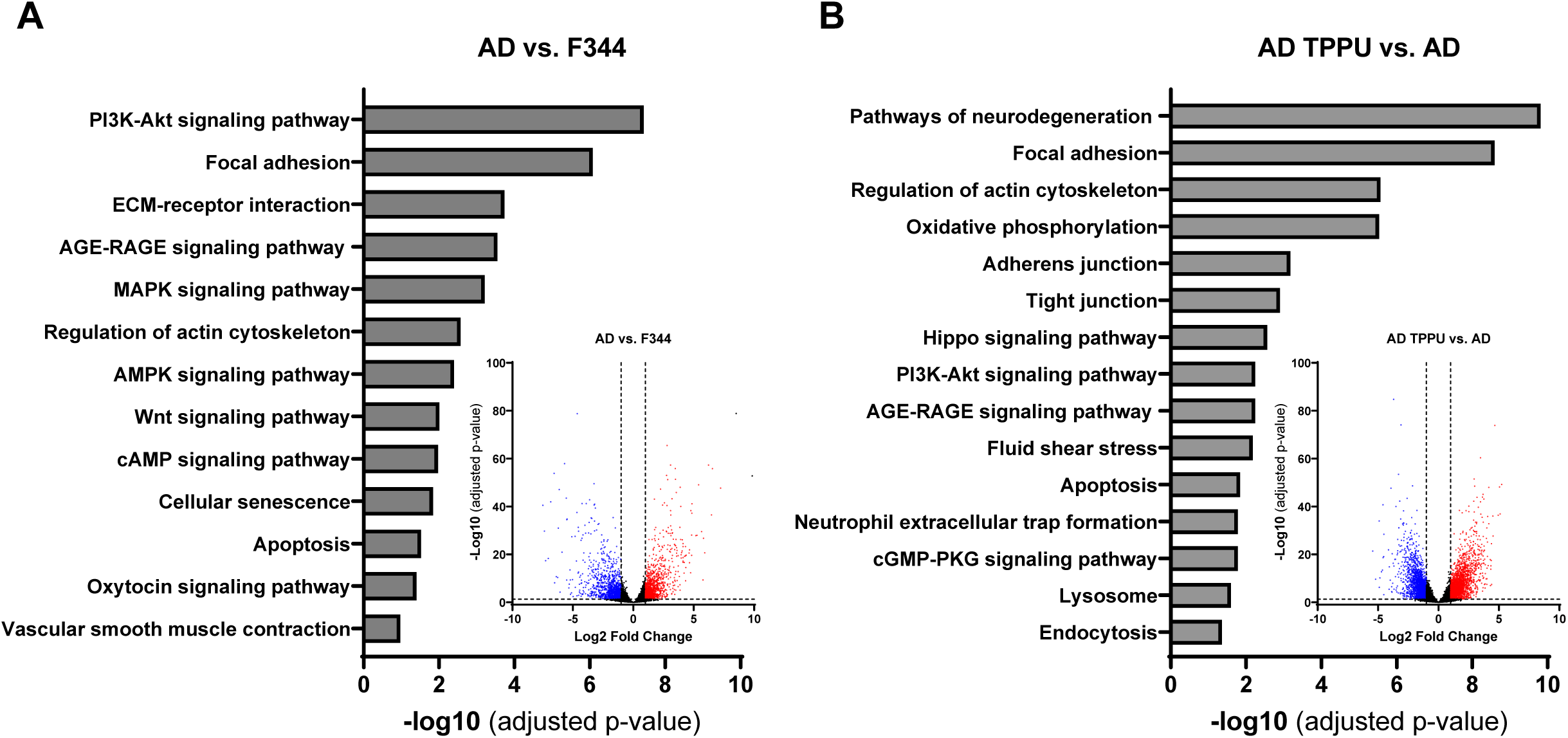
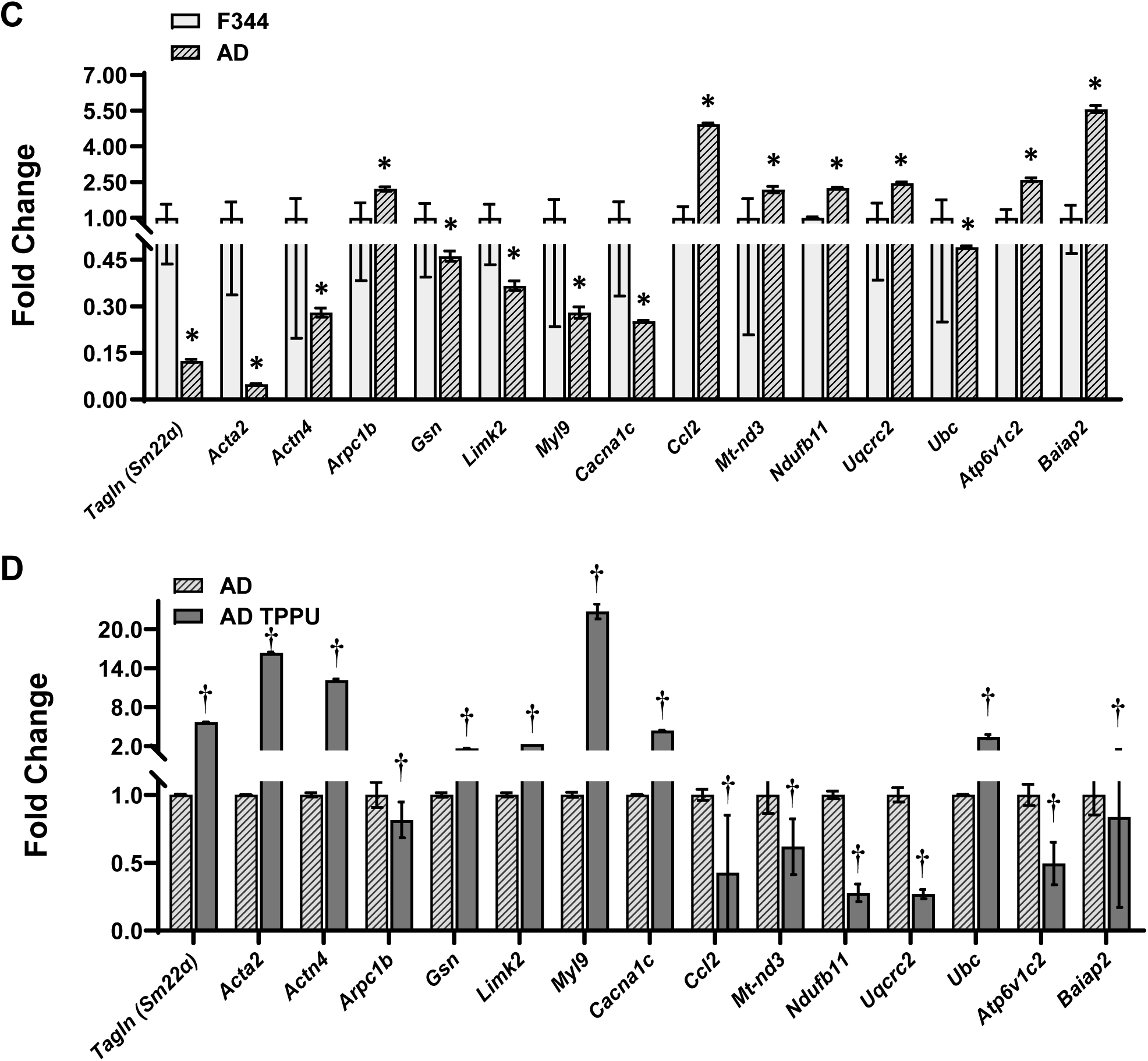
Comparison of Transcriptomic Profiling by Bulk RNA-seq in Primary Cerebral Vascular Smooth Muscle Cells (VSMCs) from TgF344-AD (AD) Rats Compared to F344 Control Rats. **A, B.** KEGG pathway enrichment analysis of the top significantly altered pathways (FDR-adjusted *p* < 0.05). Volcano plots show differential gene expression between AD and control rats. The x-axis represents the log2 fold change in gene expression, and the y-axis represents the -log10 *p*-value. Genes with |log2FC| > 1 and an FDR-adjusted *p*-value < 0.05 (or q-value) are highlighted in red (upregulated) or blue (downregulated). **C, D.** Relative gene expression levels crucial for regulating cellular contraction, oxidative stress, and inflammation. * indicates *p* < 0.05 for comparisons between cells isolated from AD rats and F344 control rats. **†** indicates *p* < 0.05 for comparisons between TPPU-treated AD cells and vehicle-treated AD cells.

### Chronic sEH Inhibition Enhances Myogenic Responses in the Cerebral Vasculature of AD and ADRD Rats

The impact of 9 weeks of sEH Inhibition on the myogenic responses of the MCA and PA was assessed by comparing age-matched DM-ADRD vs. SD rats, as well as AD vs. F344 rats. As presented in **Figure 4A**, in non-diabetic SD and F344 controls, the MCA constricted by 14.9 ± 1.6% and 16.9 ± 1.2%, respectively, when perfusion pressure elevated from 40 to 140 mmHg indicating an intact myogenic response. The myogenic response of the MCA was significantly impaired in DM-ADRD and AD animals, with vessels dilating by 40.5 ± 7.3% and 17.8 ± 5.9%, respectively, under the same conditions. Forced dilatation was observed at 120 mmHg in both DM and AD rats. However, the myogenic response of the MCAs was restored in TPPU-treated DM-ADRD and AD rats with constriction values of 22.2 ± 1.6% and 16.9 ± 3.8%, respectively, when perfusion pressure increased from 40 to 140 mmHg. In contrast, this restoration was not observed in their respective controls. Similarly, the myogenic response of the PAs was restored in TPPU-treated DM-ADRD and AD rats, but not in their respective controls (**Figure 4B**).

**Figure 4.**
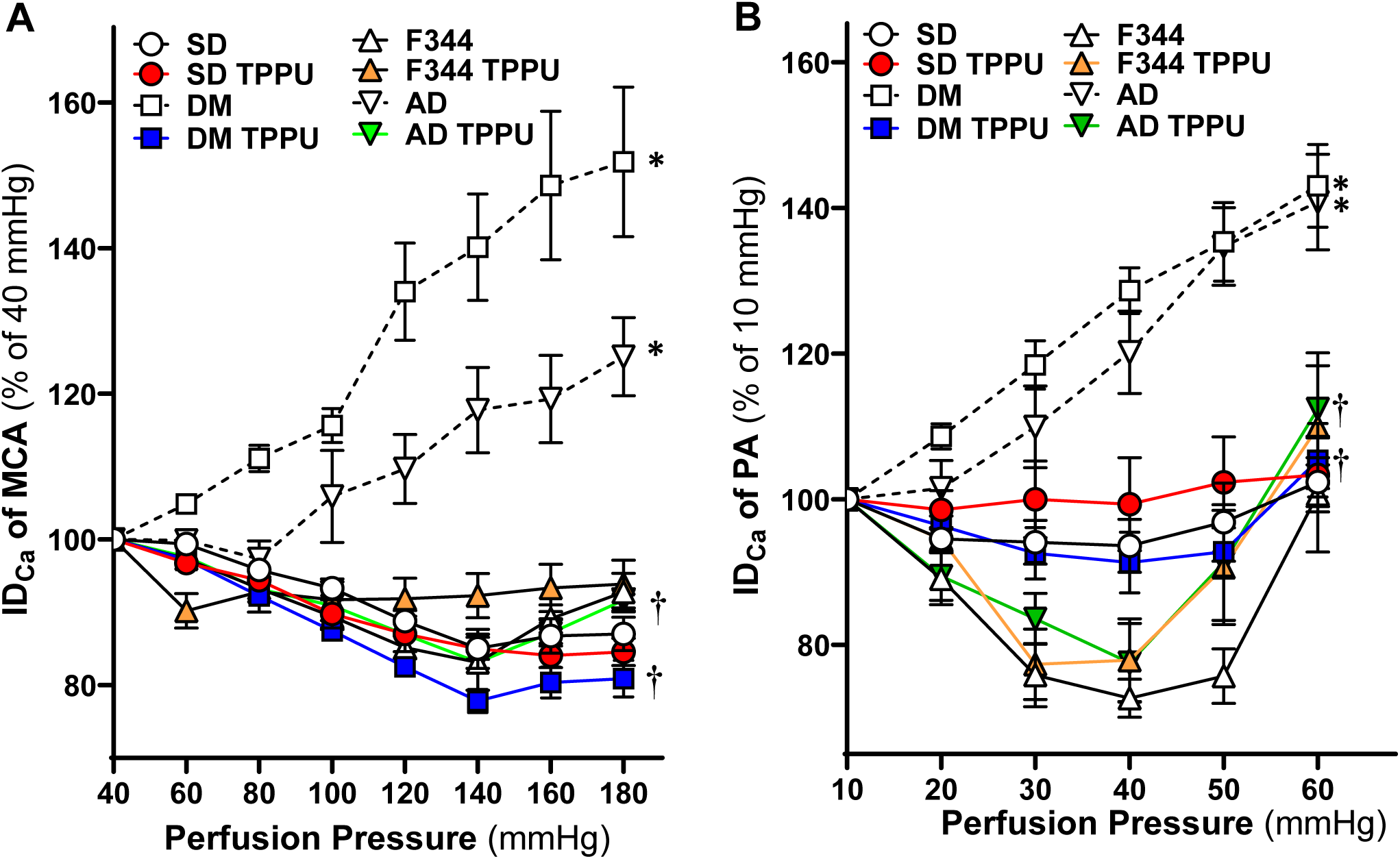
Impact of Chronic sEH Inhibition on Myogenic Responses in the Cerebral Vasculature of AD/ADRD rat. **A.** Comparison of the impact of chronic sEH inhibition on myogenic responses of middle cerebral arteries (MCAs). **B.** Comparison of the impact of chronic sEH inhibition on myogenic responses of parenchymal arterioles (PAs). Both comparisons are made between age-matched diabetic T2DN (DM) vs. Sprague Dawley (SD) rats and TgF344-AD (AD) vs. F344 rats. The rats were treated with a vehicle (1% PEG-400 in drinking water) or the sEH inhibitor TPPU (1 mg/kg/day) for 9 weeks. Mean values ± SEM are presented, with 6-16 rats per group. * denotes *p* < 0.05 for comparisons between the corresponding values in AD or DM-ADRD rats compared to their respective control rats. **†** denotes *p* < 0.05 for comparisons between the corresponding values obtained from TPPU-treated rats compared to vehicle-treated rats within the same strain.

### Acute sEH Inhibition Induces Vasodilation in the MCAs of Control Rats

We next evaluated whether TPPU directly impacts the MCAs. The IDs of the MCAs isolated from 24-36 weeks for TgF344-AD vs. F344 and 24-48 weeks for T2DN vs. SD rats were compared at 120 mmHg. As shown in **Figure 5A**, a higher dose of TPPU (10 µM) significantly induces vasodilation in SD rats within 20 minutes of treatment, whereas a low dose (1 µM) causes vasodilation starting at 30 mins. TPPU (10 µM) also induces vasodilation in the MCAs of DM-ADRD rats within 20 minutes, while TPPU (1 µM) causes MCA constriction at 30 minutes. On the other hand, high-dose TPPU dilates the MCAs in F344 rats but not in AD rats (**Figure 5B**).

**Figure 5.**
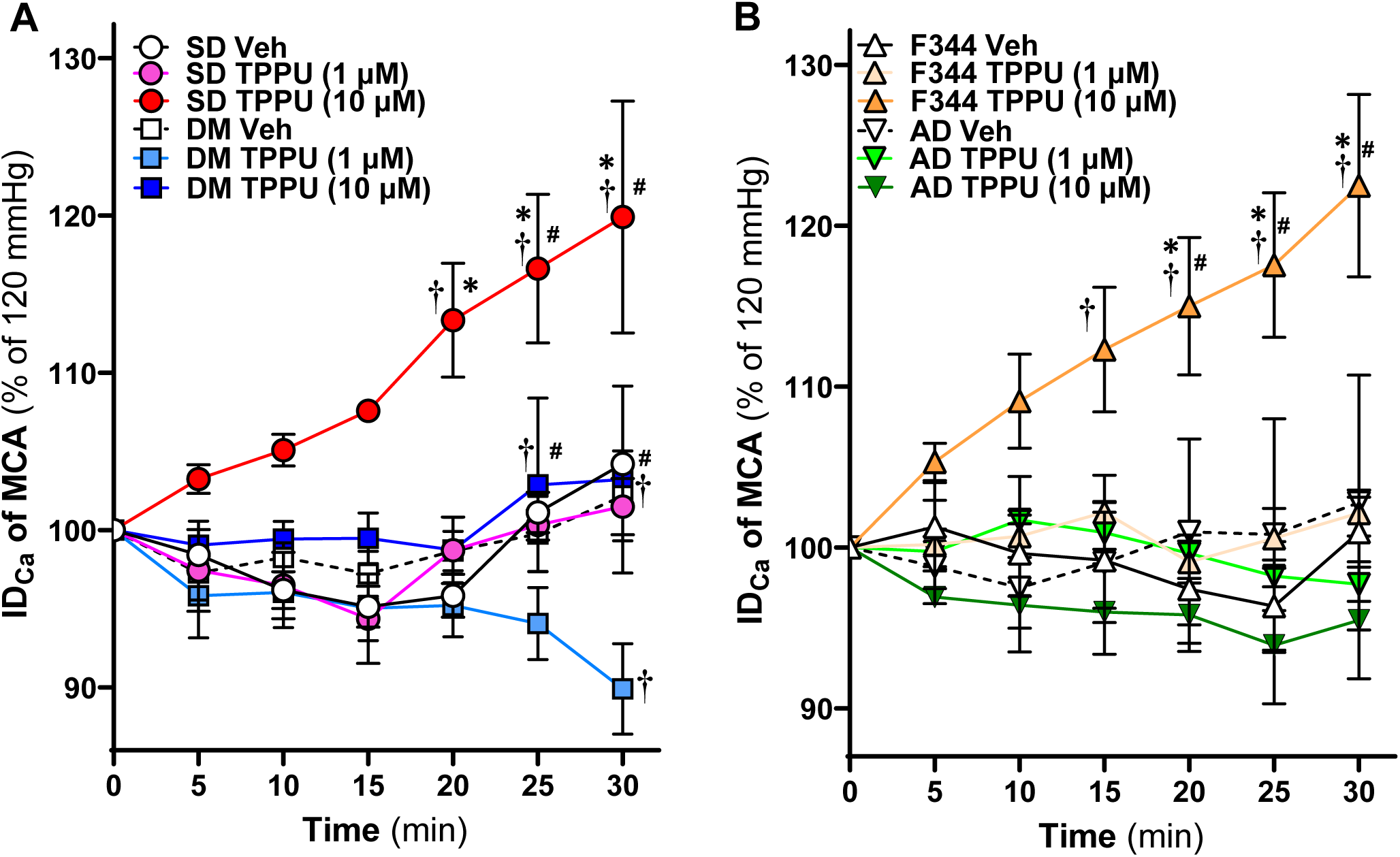
Impact of Acute sEH Inhibition on Myogenic Responses in the Cerebral Vasculature of AD/ADRD rats. **A.** Comparison of the impact of acute sEH inhibition on myogenic responses of middle cerebral arteries (MCAs) between age-matched diabetic T2DN (DM) vs. Sprague Dawley (SD) rats. **B.** Comparison of the impact of acute sEH inhibition on myogenic responses of MCAs between age-matched TgF344-AD (AD) vs. F344 rats. The vessels were treated with either a vehicle (0.1% DMOS) or the sEH inhibitor TPPU (1 and 10 µM) for 30 minutes at a perfusion pressure of 120 mmHg. Mean values ± SEM are presented, with 4-10 rats per group. * denotes *p* < 0.05 for comparisons between the corresponding values in TPPU-treated vessels at the same dosage in AD or DM-ADRD rats compared to their respective control rats. **†** denotes *p* < 0.05 for comparisons between values from TPPU-treated rats and vehicle-treated rats within the same strain. **#** denotes *p* < 0.05 for comparisons between values from TPPU (10 µM)-treated rats and TPPU (1 µM)-treated rats within the same strain.

### Chronic sEH Inhibition Enhances Acetylcholine-Induced Vasodilation in the MCA of DM-ADRD rats

Since acute sEH inhibition induces vasodilation in endothelial-intact MCAs of SD and F344 rats, the effects of TPPU on endothelial-dependent vasodilation were assessed. **Figure 6** shows that vehicle-treated MCAs in DM-ADRD rats exhibit significant endothelial dysfunction, failing to dilate to Ach. In contrast, TPPU-treated DM-ADRD rats showed dose-dependent dilation, similar to vehicle-treated SD rats.

**Figure 6.**
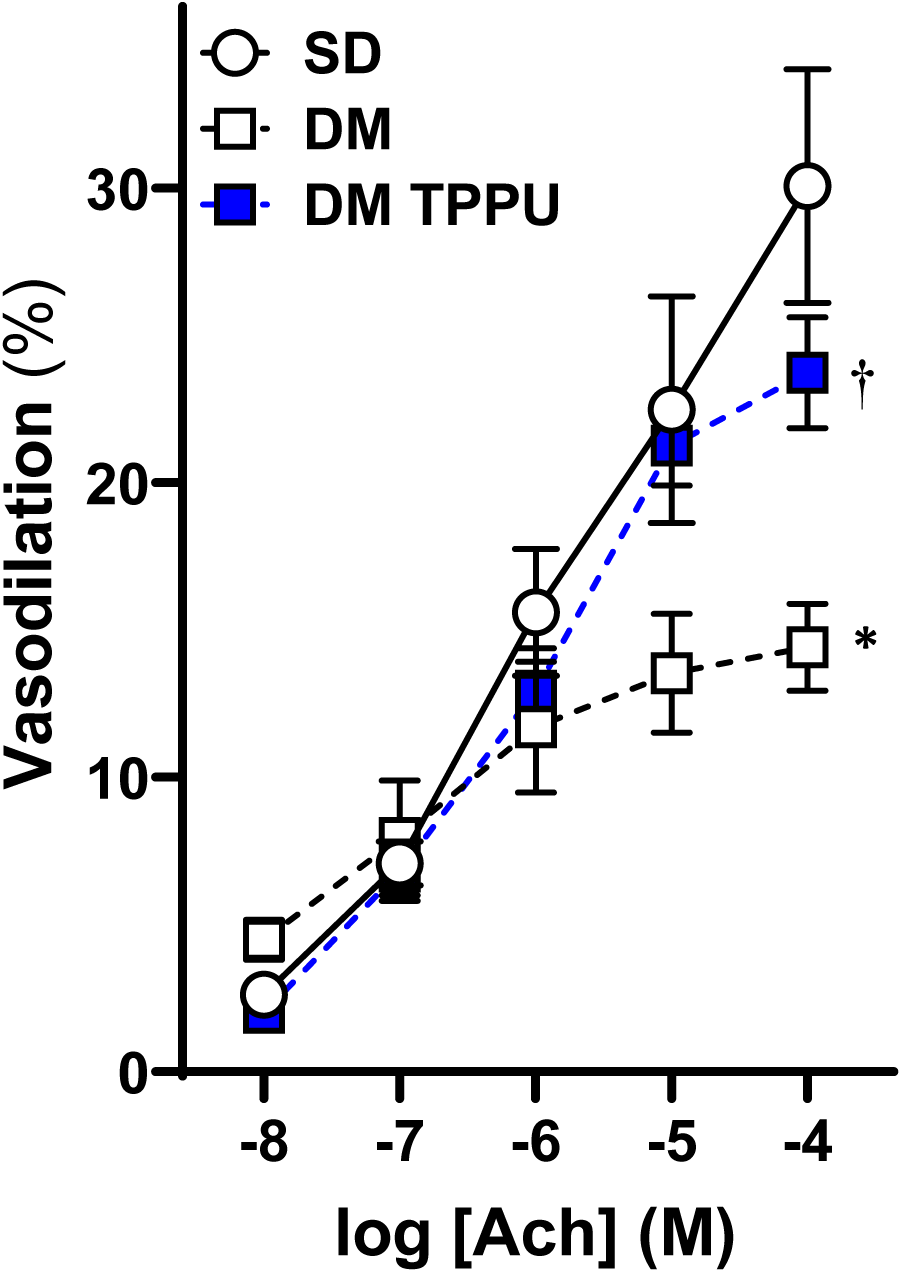
Impact of Chronic sEH Inhibition on Acetylcholine (Ach)-Induced Vasodilation in the Middle Cerebral Artery (MCA) of Diabetic-ADRD rats. Comparison of the impact of chronic sEH inhibition on Ach-induced vasodilation in the MCAs of TPPU-treated versus vehicle-treated DM-ADRD and Sprague Dawley (SD) rats. The rats were treated with a vehicle (1% PEG-400 in drinking water) or the sEH inhibitor TPPU (1 mg/kg/day) for 9 weeks. Mean values ± SEM are presented, with 4-6 rats per group. * denotes *p* < 0.05 for comparisons between the corresponding values in DM-ADRD rats and SD rats. **†** denotes *p* < 0.05 for comparisons between the corresponding values obtained from TPPU-treated rats and vehicle-treated rats within the same strain.

### Chronic sEH Inhibition Improves CBF Autoregulation of AD and ADRD Rats

The impact of 9 weeks of sEH Inhibition on CBF autoregulation was assessed by comparing age-matched DM-ADRD vs. SD rats, as well as AD vs. F344 rats using laser Doppler flowmetry with our established protocol.^17,25^ Our results align with previous findings, demonstrating that both AD and DM-ADRD rats have impaired CBF autoregulation. ^7,12,13,25,59,60^ Further, as shown in **Figure 7**, sEH inhibition improves CBF autoregulation in both AD and DM-ADRD rats. CBF increased by 11.3 ± 5.0% in vehicle-treated SD rats, 63.9 ± 9.3% in vehicle-treated DM-ADRD rats, and 27.1 ± 3.3% in TPPU-treated DM-ADRD rats when perfusion pressure rose from 100 to 140 mmHg. Conversely, when perfusion pressure decreased from 100 to 40 mmHg, the CBF decreased 19.5 ± 5.7% in vehicle-treated SD rats, 34.3 ± 4.7% in vehicle-treated DM-ADRD rats, and 33.5 ± 4.1% in TPPU-treated DM-ADRD rats. Similarly, CBF autoregulation was restored in TPPU-treated AD rats compared to vehicle-treated AD and control rats.

**Figure 7.**
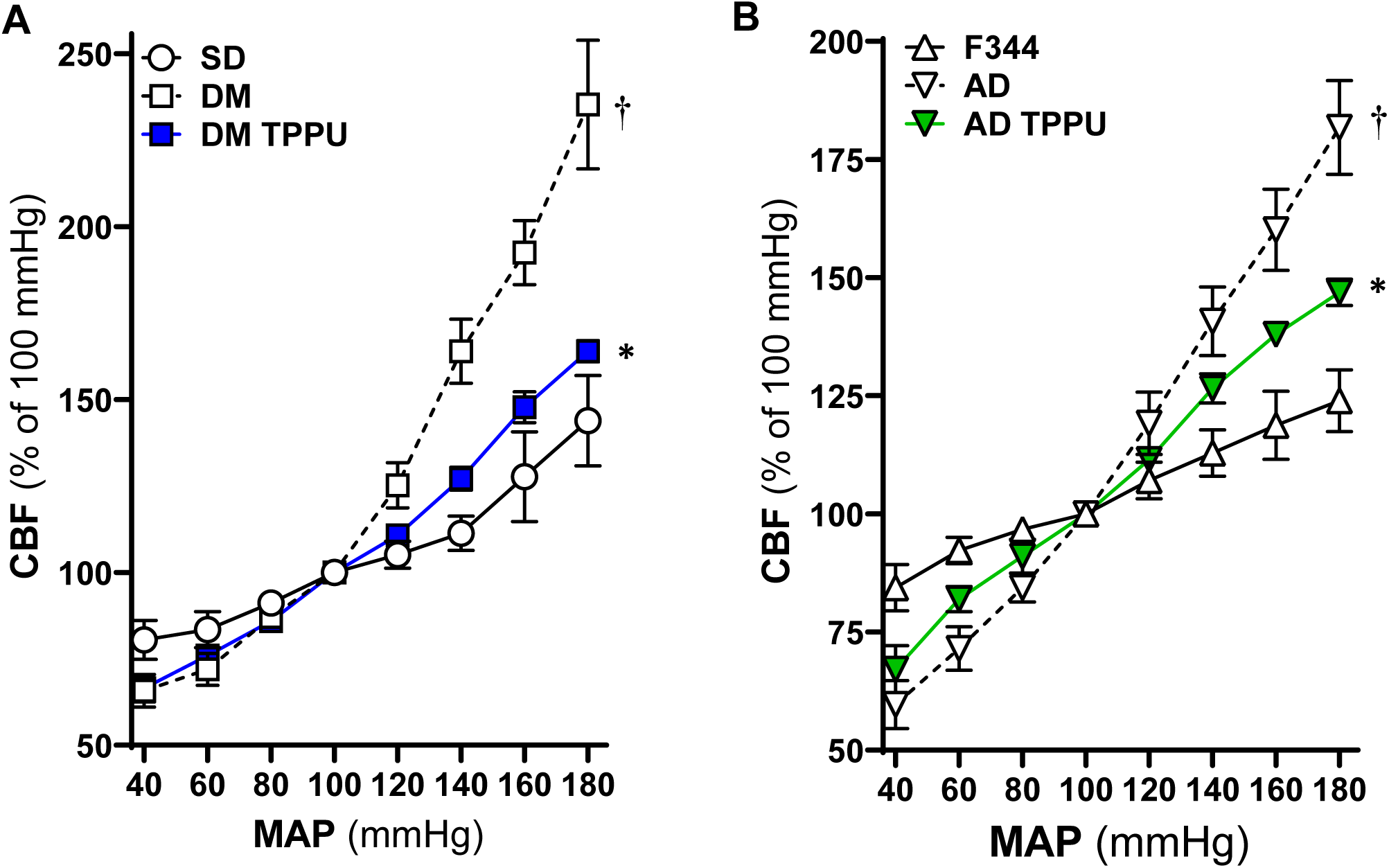
Impact of Chronic sEH Inhibition on Autoregulation of the Cerebral Blood Flow (CBF) in AD and Diabetic-ADRD rats. **A.** Comparison of the impact of chronic sEH inhibition on CBF autoregulation in TPPU-treated versus vehicle-treated DM-ADRD and Sprague Dawley (SD) rats. **B.** Comparison of the impact of chronic sEH inhibition on CBF autoregulation in TPPU-treated versus vehicle-treated AD and F344 rats. The rats were treated with a vehicle (1% PEG-400 in drinking water) or the sEH inhibitor TPPU (1 mg/kg/day) for 9 weeks. Mean values ± SEM are presented, with 4-10 rats per group. * denotes *p* < 0.05 for comparisons between the corresponding values in AD rats compared to F344 rats. **†** denotes *p* < 0.05 for comparisons between the corresponding values obtained from TPPU-treated rats and vehicle-treated rats within the same strain.

### Effects of Inhibition of sEH on Contractile Capability of Cerebral VSMCs Isolated from AD and F344 Rats

The results from acute and chronic sEH inhibition indicate that TPPU may enhance VSMC-dependent myogenic responses and autoregulation at low doses, as well as endothelial-dependent vasodilation at high doses. To validate this finding, the effect of low-dose TPPU was evaluated on the contractile capacity of VSMCs of AD compared to control rats. The optimal dosages (0.1, 1, and 10 µM) and duration of the treatment were first examined using cerebral VSMCs isolated from AD rats (data not shown). As depicted in **Figure 8**, TPPU-treated AD cells display enhanced contraction compared to untreated or vehicle-treated AD cells.

**Figure 8.**
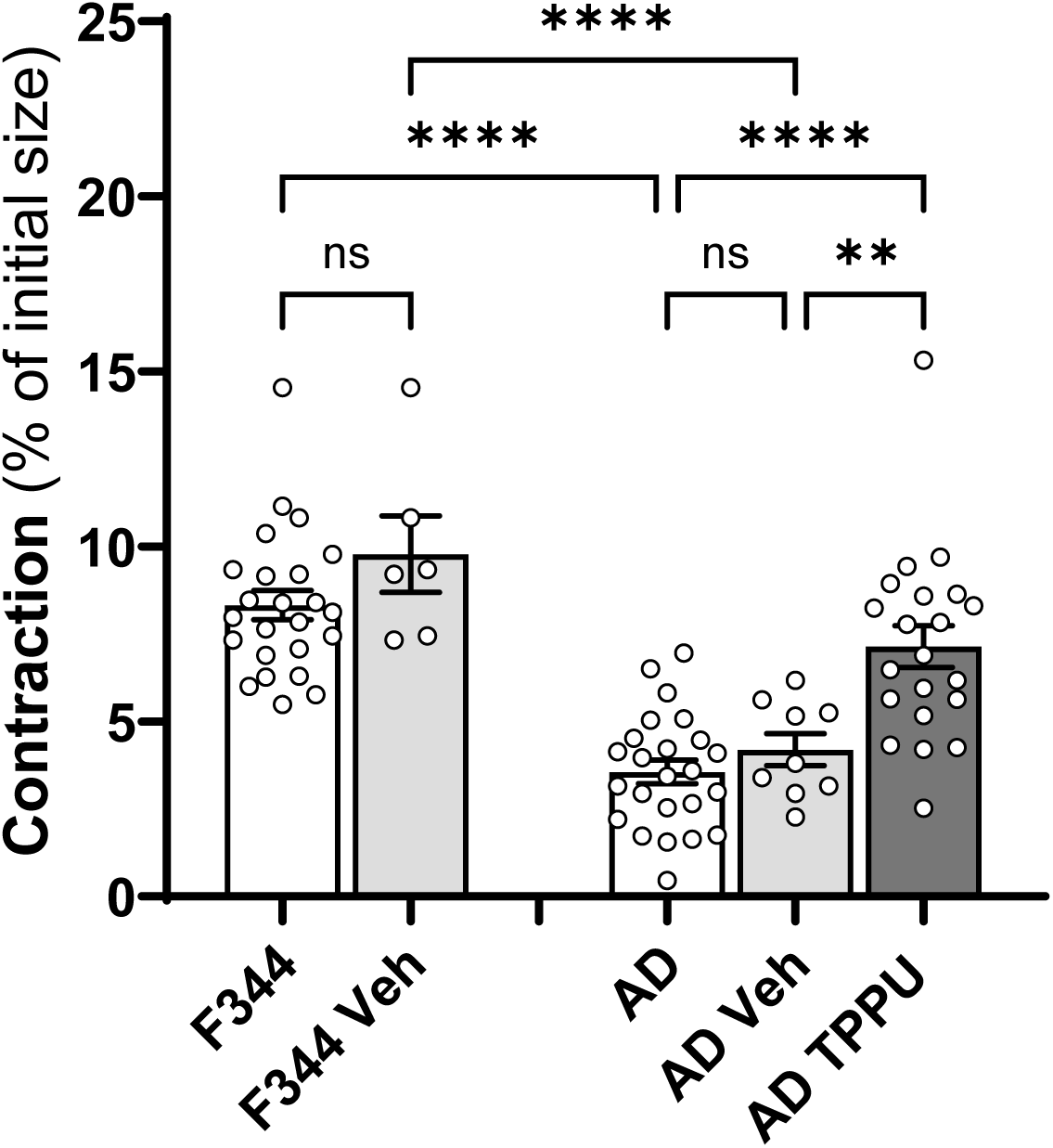
Impact of Chronic sEH Inhibition on Contractile Capability of Primary Cerebral Vascular Smooth Muscle Cells (VSMCs) Isolated from F344 and AD rats. Comparison of the impact of chronic sEH inhibition on contractile capability on cerebral VSMCs of AD and F344 rats. Mean values ± SEM are presented, with 3 rats, and experiments were repeated three times per group. ** denotes *p* < 0.01 and **** denotes *p* < 0.0001 for comparisons between the indicated values and their respective controls.

### Chronic sEH Inhibition Diminishes the Production of ROS and Mitochondrial Superoxide in the MCA of AD and ADRD Rats

As shown in **Figure 9**, the fold change in DHE fluorescence intensity was significantly higher in freshly isolated MCAs from DM-ADRD (8.58 ± 1.36, n = 4) and AD (6.71 ± 0.40, n = 5) rats compared to control SD (1.00 ± 0.08, n = 5) and F344 rats (1.00 ± 0.05, n = 6). TPPU treatment reduced DHE intensity in DM-ADRD (2.02 ± 0.21, n = 17) and AD (0.83 ± 0.09, n = 6) vessels to statistical control levels, specifically. Similarly, as shown in **Figure 10**, the fold change in MitoSOX fluorescence intensity was higher in DM-ADRD (3.84 ± 0.18, n = 8) and AD (2.77 ± 0.44, n = 4) rats compared to SD (1.00 ± 0.32, n = 4) and F344 rats (1.00 ± 0.13, n = 6). TPPU treatment reduced MitoSOX fluorescence intensity in DM-ADRD (1.49 ± 0.18, n = 13) and AD (1.42 ± 0.28, n = 11) vessels to statistical control levels, specifically.

**Figure 9.**
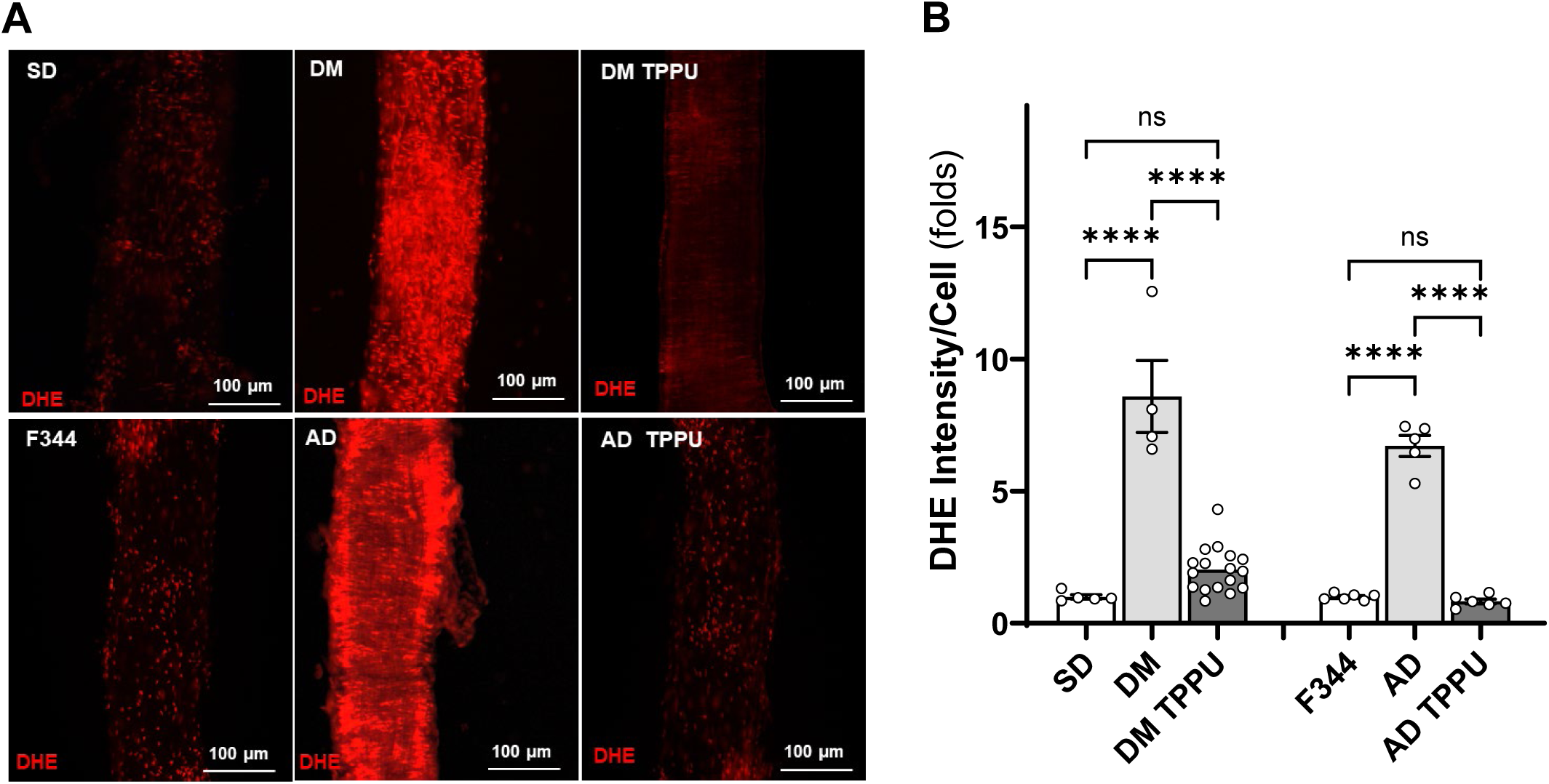
Impact of Chronic sEH Inhibition on the Production of Reactive Oxygen Species (ROS) in the MCA of AD and ADRD Rats. **A.** Representative images of DHE staining in MCA freshly isolated from age-matched diabetic T2DN (DM) vs. Sprague Dawley (SD) rats and TgF344-AD (AD) vs. F344 rats. The rats were treated with either a vehicle (1% PEG-400 in drinking water) or the sEH inhibitor TPPU (1 mg/kg/day) for 9 weeks. **B.** Quantitative analysis of the fold change of DHE fluorescence. Mean values ± SEM are presented, with 4 -17 rats per group. **** denotes *p* < 0.0001 for comparisons between the indicated values and their respective controls.

**Figure 10.**
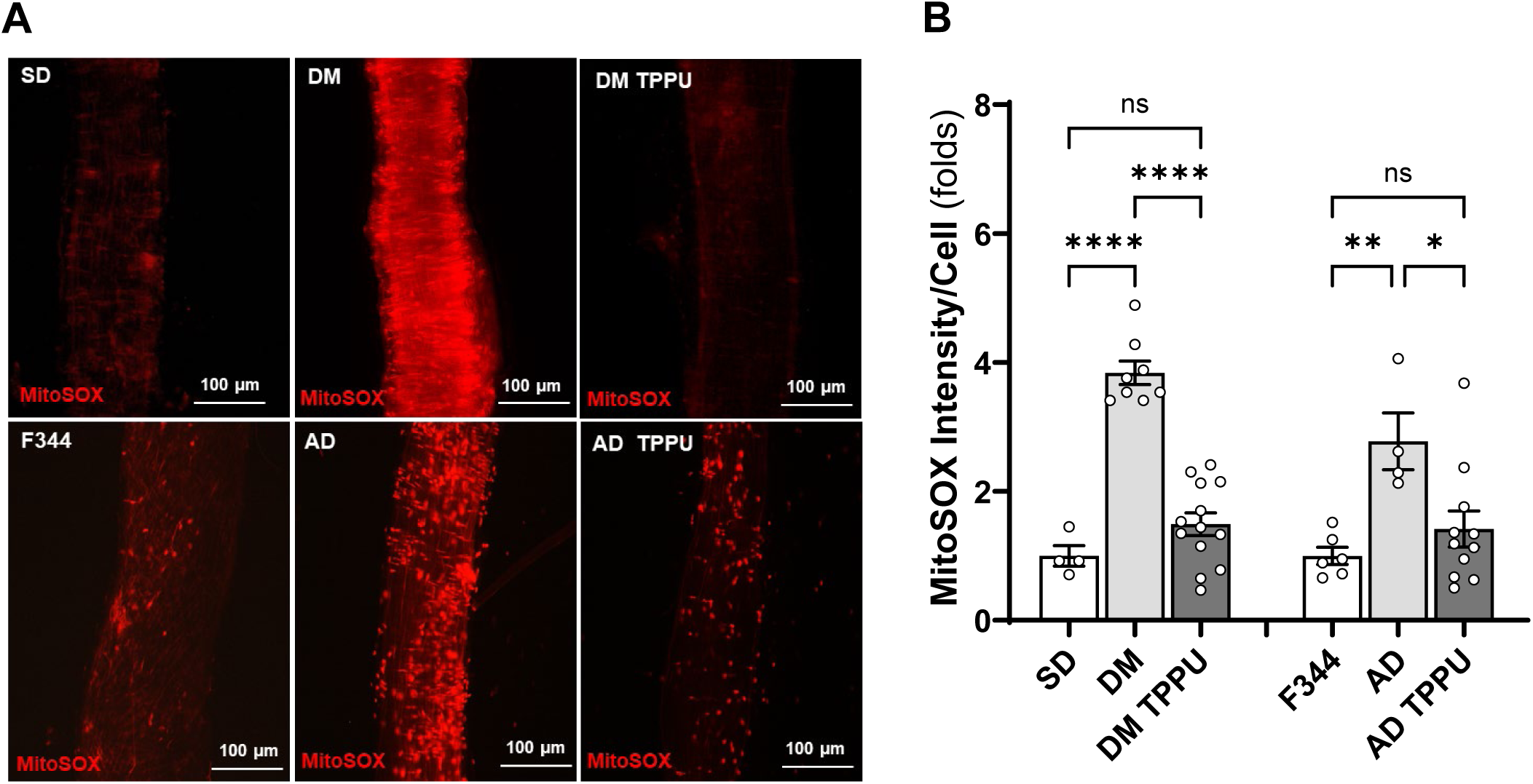
Impact of Chronic sEH Inhibition on the Production of Mitochondrial Superoxide in the MCA of AD and ADRD Rats. **A.** Representative images of MitoSOX staining in MCA freshly isolated from age-matched diabetic T2DN (DM) vs. Sprague Dawley (SD) rats and TgF344-AD (AD) vs. F344 rats. The rats were treated with a vehicle (1% PEG-400 in drinking water) or the sEH inhibitor TPPU (1 mg/kg/day) for 9 weeks. **B.** Quantitative analysis of the fold change of MitoSOX fluorescence. Mean values ± SEM are presented, with 4 -13 rats per group. * denotes *p* < 0.05, ** denotes *p* < 0.01, and **** denotes *p* < 0.0001 for comparisons between the indicated values and their respective controls.

## DISCUSSION

AD/ADRD poses a major public health challenge, with a significant rise in cases predicted due to an aging population. Millions of individuals over 65 are affected in the U.S., with numbers expected to double by 2060. In 2023, AD/ADRD costs totaled $345 billion in the U.S., and global dementia care costs amount to $1.3 trillion annually.^1^ Kisunla™ (donanemab-azbt) received FDA approval in July 2024 for the treatment of early symptoms of AD.^61^ It is the third FDA-approved drug targeting Aβ, joining aducanumab and lecanemab.^3^ Despite these advancements, amyloid-targeting therapies are associated with side effects such as amyloid-related imaging abnormalities, severe headaches, and anaphylaxis.^62^ Furthermore, the long-term effectiveness of these anti-amyloid drugs remains to be fully established. Overall, there is still no effective cure for AD/ADRD.

The most prevalent form of dementia involves a comorbidity of AD and vascular disease.^63^ DM is a major cardiovascular risk factor for ADRD.^64^ DM-ADRD shares several pathological features with AD, characterized by a complex interplay of mechanisms, including the accumulation of Aβ plaques, tau tangles, and neuroinflammation.^65^ Reduced CBF significantly contributes to AD/ADRD development, with cerebrovascular dysfunction like impaired CBF autoregulation, BBB damage, and neurovascular uncoupling leading to brain hypoperfusion. This hypoperfusion often precedes Aβ and tau abnormalities, initiating neuronal damage. Inflammation related to cerebrovascular dysfunction also plays a crucial role, and brain hypoperfusion worsens cognitive decline, particularly in individuals with risk factors such as hypertension and diabetes.

EETs, the substates of sEH in the AA pathway, reduce inflammation and act as endothelial-derived hyperpolarizing factors until they are converted to DHETs by the sEH.^66^ DiHOMEs, the metabolites of sEH in the LA pathway, function as both markers and contributors to tissue inflammation in several diseases.^67,68^ Recent animal studies have shown that inhibiting sEH protects various conditions, ^29–37^ including cognitive impairments. ^38,39^ Genome-wide studies have identified associations between single nucleotide polymorphisms in the sEH-encoding gene *EPHX2* and AD/ADRD. ^69–73^ Genetic deletion ^70^ or inhibition of sEH improved cognition in AD ^40,69,74^ and DM-related ADRD. ^75–77^ While most of these studies showed that cognitive protection by sEH inhibition was due to its anti-inflammatory and neuronal protective effects, its vascular contribution to AD/ADRD has been largely unknown.

In the present study, the effects of sEH inhibition with TPPU, which has cognitive protection due to its anti-inflammatory and neuronal protective effects,^40,69,74^ were assessed on cerebral vascular function and cognition in rat models of AD and DM-related ADRD. First, we evaluated the impact of TPPU treatment on body weight, glucose levels, and HbA1C. We found that 9 weeks of TPPU treatment did not affect body weight but significantly improved cognitive function in AD and DM-ADRD rats. Notably, reductions in plasma glucose and HbA1C levels were observed only in DM-ADRD rats. Luo et al. found that sEH inhibition lowered blood glucose levels in obese mice fed a high-fat, high-salt diet but not in non-obese mice.^78^ This effect was linked to improved insulin resistance, reduced hypertension, and decreased renal SGLT2 expression, attributed to the inhibition of the IκB kinase α/β/NF-κB inflammatory pathway in obese mice. While sEH inhibition has been shown to improve glucose homeostasis and insulin resistance in other obese-diabetes models,^79,80^ some studies report inconsistencies, potentially due to variations in compounds and treatment durations. ^81^

Chronic sEH inhibition enhances the myogenic response and CBF autoregulation in the cerebral circulation of both AD and DM-ADRD rats. The myogenic response is an intrinsic property of VSMCs that regulates CBF by causing constriction of cerebral arteries and arterioles in response to increases in blood pressure. ^59^ These reactivities play a crucial role in regulating brain perfusion and maintaining stable blood perfusion to the brain, which is essential for normal brain function. When blood pressure increases, cerebral arteries can constrict to prevent excessive blood flow and maintain stable perfusion pressure. Conversely, when blood pressure decreases, these arteries can dilate to increase blood flow and ensure adequate oxygen and nutrient delivery to brain tissue. We previously observed reduced contractile capabilities in cerebral VSMCs in both AD^7^ and DM-ADRD,^22,43^ with Aβ directly impairing VSMC constriction.^7^ Additionally, we reported elevated ROS and mitochondrial superoxide production in primary cerebral VSMCs and freshly isolated cerebral vessels from AD and DM-ADRD rats, which impair their contractile capabilities.^7,22,25^ Thus, our results afford a mechanistic hypothesis that the observed benefits of chronic sEH inhibition might be due to reduced ROS and mitochondrial superoxide levels, as indicated by decreased DHE and MitoSOX staining in the MCAs of AD and DM-ADRD rats. From this perspective, it appears that sEH inhibition may enhance cerebral VSMC function by mitigating oxidative stress. This observation was validated by assessing the contractile function and transcriptomic profiling of primary VSMCs isolated from AD rats treated with TPPU. We found that AD cells treated with TPPU exhibited improved contraction compared to untreated or vehicle-treated AD cells. Bulk RNA-seq results revealed that, in AD VSMCs, the expression of genes regulating cellular contraction was lower, while genes associated with oxidative stress and inflammation were higher. TPPU treatment reversed these changes. Among these genes, *Tagln* or *Sm22*α (transgelin),^82^ *Acta2* (α-SMA),^21^ *Actn4* (ACTN4),^83^ *Arpc1b* (ARP2/3 1B),^84^ *Gsn* (GSN),^85^ *Limk2* (LIMK2),^86^ *Myl9* (MYL9),^87^ *Cacna1c* (CACIC)^88^ play a crucial role in regulating smooth muscle cell contraction and cytoskeletal organization. *Ccl2* (MCP1),^89^ *Mt-nd3* (ND3),^90^ *Ndufb11* (NDUFB11),^91^ *Uqcrc2* (UQCRC2),^92^ and *Ubc* (Ubiquitin C)^93^ are involved in inflammation and oxidative stress, which may affect VSMC function in AD/ADRD. Additionally, *Atp6v1c2* (ATP6V1C2), which regulates intracellular pH balance, and *Baiap2* (BAIP2), which influences endothelial and VSMC dynamics, also showed significant alterations.

Acute sEH inhibition induces vasodilation in the MCAs of SD rats, with a higher dose of TPPU (10 µM) showing effects within 20 minutes, and a lower dose (1 µM) taking 30 minutes to show its effects. Interestingly, TPPU (10 µM) also induces vasodilation in the MCAs of DM-ADRD rats within 20 minutes, while TPPU (1 µM) causes MCA constriction at 30 minutes. High-dose TPPU dilates the MCAs in F344 rats but not in AD rats. Furthermore, chronic sEH inhibition enhances Ach-induced vasodilation in the MCA of DM-ADRD rats. These results suggest that lower doses or shorter durations of sEH inhibition enhance VSMC function, while higher doses or longer durations are required to fully achieve the endothelial-derived hyperpolarizing effects of EETs. Although our study alone does not sufficiently make this conclusion, it aligns with findings by Ostermann et al.,^41^ who observed a steady-state blood concentration of ∼0.9 µM TPPU after 8 days of oral treatment at 0.5 mg/kg/day in SD rats. Based on this, we speculate that the relatively “low” TPPU levels achieved with our 9-week treatment at 1 mg/kg/day could explain the observed enhancement in VSMC function.

Notably, in this study, key experiments on vascular and cognitive functions were conducted at 15 months of age for DM-ADRD rats and 9 months of age for AD rats, compared to age-matched controls. In contrast, primary VSMCs were isolated from 3-week-old AD and F344 rats. The TgF344-AD rat model, which is genetically modified to overexpress mutated human amyloid precursor protein (APP) with the Swedish mutation and human presenilin 1 (PS1) with the Δ exon 9 mutation on an F344 background, exhibits age-dependent increases in Aβ levels^94^ and is thus expected to show elevated Aβ expression even at a young age. We have reported that Aβ (1-42) reduced cerebral cell contraction of primary VSMCs isolated from 3-week-old F344 rats, similar to those from AD rats at the same age. ROS production and mitochondrial ROS were enhanced, and mitochondrial respiration and ATP production were reduced in AD VSMCs compared with F344 cells.^7^ However, in this AD model, vascular and cognitive dysfunctions were observed at 16 and 24 weeks, respectively,^24,94^ suggesting that disease progression is gradual, even with genetic predispositions related to Aβ. This gradual progression might explain the lack of significant changes in EPHX2 expression observed in our bulk RNA-seq studies, despite a trend indicating higher levels in AD versus F344 and AD-TPPU groups. Nevertheless, the findings from our cell contraction assays and bulk RNA-seq are consistent with the observed vascular and cognitive function outcomes. Future studies will focus on analyzing primary VSMCs isolated from TPPU-treated AD rats at the age when these functional studies are performed.

We acknowledge several limitations of the present study. Notably, we did not assess the impact of sEH inhibition on blood pressure, even though our previous research indicated no significant baseline differences in blood pressure between AD/ADRD models and their respective controls.^7,25,43^ This omission is relevant because increased levels of EETs, which may induce vasodilation and lower blood pressure, have been observed in both animal and human studies.^95^ Additionally, we could not determine whether the vascular and cognitive benefits of TPPU in DM-ADRD rats are directly attributable to sEH inhibition or if they result from downstream effects of reduced plasma glucose and HbA1c levels, as hyperglycemia itself can damage cerebral vascular cells.^11,22,23,96^ While we validated the contractile function of primary cerebral VSMCs and performed oxidative stress staining using DHE and MitoSOX on isolated vessels, we did not assess the differential mRNA expression of genes identified through bulk RNA-seq in response to sEH inhibition. Future research should also explore drug concentration and BBB permeability to further clarify our findings. Despite these limitations, our study provides valuable insights into the effects of sEH inhibition on VSMC function and cerebral vascular health in AD and DM-ADRD models, highlighting both its potential benefits and the need for further investigation to refine treatment strategies. For instance, while approximately 20% of TPPU crosses the BBB in AD mice,^33,40^ modifying TPPU to reduce its BBB permeability could decrease neuronal toxicity^97^ and enhance its benefits for endothelial cells and VSMCs. Furthermore, our findings could inform the use of EC5026,^37^ a compound structurally similar to TPPU but with higher potency for human sEH enzymes. EC5026 has completed several clinical trials, including two Phase 1a studies (NCT04228302 and NCT04908995) and a Phase 1b Multiple Ascending Dose study (NCT06089837), confirming its safety, tolerability, and pharmacokinetics in healthy volunteers. Given its efficacy in rodent models and the absence of notable side effects in humans, EC5026 is expected to offer comparable benefits for vascular, neuronal, and cognitive functions in AD/ADRD patients, positioning it as a promising candidate for future clinical trials.

## DECLARATIONS

### Author Contributions

F.F. conceived and designed research; C.T., J.J.B., H.Z., A.G., X.F., Y.L., S.W., S. H.H., W.G., G.C.M., J.S., D.B., and C.C. performed experiments; C.T., J.J.B., H.Z. A.G., S.B., G.C.M. and F.F. analyzed data; C.T., J.J.B., H.Z. A.G., P.O’H., Z.B., J.A.F., Y.D., H.Y., B.D.H., R.J.R., and F.F. interpreted results; C.T., A.G., and F.F. prepared figures; F.F. drafted the manuscript; C.T., J.J.B., H.Z., A.G., S.B., X.F., Y.L., S.W., S.H.H., W.G., G.C.M., J.S., D.B., C.C., K.W., C.M., J.Y., S.M.S., P.O’H., Z.B., J.A.F., Y.D., H.Y., B.D.H., R.J.R., and F.F. edited and revised the manuscript; all authors approved the final version of the manuscript.

## REFERENCES

1. 2023 Alzheimer’s disease facts and figures. Alzheimers Dement. 2023;19:1598–1695. doi: 10.1002/alz.13016

2. Huang LK, Chao SP, Hu CJ. Clinical trials of new drugs for Alzheimer disease. J Biomed Sci. 2020;27:18. doi: 10.1186/s12929-019-0609-7

3. Fang X, Zhang J, Roman RJ, Fan F. From 1901 to 2022, how far are we from truly understanding the pathogenesis of age-related dementia? GeroScience. 2022;44:1879–1883. doi: 10.1007/s11357-022-00591-7

4. Bracko O, Cruz Hernández JC, Park L, Nishimura N, Schaffer CB. Causes and consequences of baseline cerebral blood flow reductions in Alzheimer’s disease. J Cereb Blood Flow Metab. 2021;41:1501–1516. doi: 10.1177/0271678x20982383

5. Korte N, Nortley R, Attwell D. Cerebral blood flow decrease as an early pathological mechanism in Alzheimer’s disease. Acta Neuropathologica. 2020;140:793–810. doi: 10.1007/s00401-020-02215-w

6. Fan F, Roman RJ. Reversal of cerebral hypoperfusion: a novel therapeutic target for the treatment of AD/ADRD? Geroscience. 2021;43:1065–1067. doi: 10.1007/s11357-021-00357-7

7. Fang X, Tang C, Zhang H, Border JJ, Liu Y, Shin SM, Yu H, Roman RJ, Fan F. Longitudinal characterization of cerebral hemodynamics in the TgF344-AD rat model of Alzheimer’s disease. Geroscience. 2023. doi: 10.1007/s11357-02300773-x

8. Fan F, Pabbidi M, Lin RCS, Ge Y, Gomez-Sanchez EP, Rajkowska GK, Moulana M, Gonzalez-fernandez E, Sims J, Elliott MR, et al. Impaired myogenic response of MCA elevates transmission of pressure to penetrating arterioles and contributes to cerebral vascular disease in aging hypertensive FHH rats. The FASEB Journal. 2016;30:953.957–953.957. doi: 10.1096/fasebj.30.1_supplement.953.7

9. Fan F, Ge Y, Lv W, Elliott MR, Muroya Y, Hirata T, Booz GW, Roman RJ. Molecular mechanisms and cell signaling of 20-hydroxyeicosatetraenoic acid in vascular pathophysiology. Front Biosci (Landmark Ed). 2016;21:1427–1463.

10. Crumpler R, Roman RJ, Fan F. Capillary Stalling: A Mechanism of Decreased Cerebral Blood Flow in AD/ADRD. J Exp Neurol. 2021;2:149–153. doi: 10.33696/neurol.2.048

11. Fan F, Booz GW, Roman RJ. Aging diabetes, deconstructing the cerebrovascular wall. Aging (Albany NY). 2021;13:9158–9159. doi: 10.18632/aging.202963

12. Wang S, Tang C, Liu Y, Border JJ, Roman RJ, Fan F. Impact of impaired cerebral blood flow autoregulation on cognitive impairment. Frontiers in Aging. 2022;3. doi: 10.3389/fragi.2022.1077302

13. Fang X, Crumpler RF, Thomas KN, Mazique JN, Roman RJ, Fan F. Contribution of cerebral microvascular mechanisms to age-related cognitive impairment and dementia. Physiol Int. 2022. doi: 10.1556/2060.2022.00020

14. Kinney JW, Bemiller SM, Murtishaw AS, Leisgang AM, Salazar AM, Lamb BT. Inflammation as a central mechanism in Alzheimer’s disease. Alzheimers Dement (N Y). 2018;4:575–590. doi: 10.1016/j.trci.2018.06.014

15. Finger CE, Moreno-Gonzalez I, Gutierrez A, Moruno-Manchon JF, McCullough LD. Age-related immune alterations and cerebrovascular inflammation. Molecular Psychiatry. 2022;27:803–818. doi: 10.1038/s41380-021-01361-1

16. Wang S, Jiao F, Guo Y, Booz G, Roman R, Fan F. Role of Vascular Smooth Muscle Cells in Diabetes-related Vascular Cognitive Impairment. Stroke. 2019;50:ATP556–ATP556.

17. Wang S, Zhang H, Liu Y, Li L, Guo Y, Jiao F, Fang X, Jefferson JR, Li M, Gao W, et al. Sex differences in the structure and function of rat middle cerebral arteries. Am J Physiol Heart Circ Physiol. 2020;318:H1219–H1232. doi: 10.1152/ajpheart.00722.2019

18. Pabbidi MR, Mazur O, Fan F, Farley JM, Gebremedhinm D, Harder DR, Roman RJ. Enhanced large conductance K+ channel (BK) activity contributes to the impaired myogenic response in the cerebral vasculature of Fawn Hooded Hypertensive rats. Am J Physiol Heart Circ Physiol. 2014;2014 Apr 1;306(7):H989–H1000.

19. Aguilar-Pineda JA, Vera-Lopez KJ, Shrivastava P, Chávez-Fumagalli MA, Nieto-Montesinos R, Alvarez-Fernandez KL, Goyzueta Mamani LD, Davila Del-Carpio G, Gomez-Valdez B, Miller CL, et al. Vascular smooth muscle cell dysfunction contribute to neuroinflammation and Tau hyperphosphorylation in Alzheimer disease. iScience. 2021;24:102993. doi: 10.1016/j.isci.2021.102993

20. Xu X, Liu XQ, Liu XL, Wang X, Zhang WD, Huang XF, Jia FY, Kong P, Han M. SM22α Deletion Contributes to Neurocognitive Impairment in Mice through Modulating Vascular Smooth Muscle Cell Phenotypes. Int J Mol Sci. 2023;24. doi: 10.3390/ijms24087117

21. Liu Y, Zhang H, Wu CY, Yu T, Fang X, Ryu JJ, Zheng B, Chen Z, Roman RJ, Fan F. 20-HETE-promoted cerebral blood flow autoregulation is associated with enhanced pericyte contractility. Prostaglandins Other Lipid Mediat. 2021;154:106548. doi: 10.1016/j.prostaglandins.2021.106548

22. Guo Y, Wang S, Liu Y, Fan L, Booz GW, Roman RJ, Chen Z, Fan F. Accelerated cerebral vascular injury in diabetes is associated with vascular smooth muscle cell dysfunction. Geroscience. 2020;42:547–561. doi: 10.1007/s11357-02000179-z

23. Liu Y, Zhang H, Wang S, Guo Y, Fang X, Zheng B, Gao W, Yu H, Chen Z, Roman RJ, et al. Reduced pericyte and tight junction coverage in old diabetic rats are associated with hyperglycemia-induced cerebrovascular pericyte dysfunction. Am J Physiol Heart Circ Physiol. 2021;320:H549–H562. doi: 10.1152/ajpheart.00726.2020

24. Fang X, Border JJ, Rivers PL, Zhang H, Williams JM, Fan F, Roman RJ. Amyloid beta accumulation in TgF344-AD rats is associated with reduced cerebral capillary endothelial Kir2.1 expression and neurovascular uncoupling. Geroscience. 2023. doi: 10.1007/s11357-023-00841-2

25. Wang S, Lv W, Zhang H, Liu Y, Li L, Jefferson JR, Guo Y, Li M, Gao W, Fang X, et al. Aging exacerbates impairments of cerebral blood flow autoregulation and cognition in diabetic rats. Geroscience. 2020;42:1387–1410. doi: 10.1007/s11357-020-00233-w

26. Morisseau C, Hammock BD. Impact of soluble epoxide hydrolase and epoxyeicosanoids on human health. Annu Rev Pharmacol Toxicol. 2013;53:37–58. doi: 10.1146/annurev-pharmtox-011112-140244

27. Sura P, Sura R, Enayetallah AE, Grant DF. Distribution and expression of soluble epoxide hydrolase in human brain. J Histochem Cytochem. 2008;56:551–559. doi: 10.1369/jhc.2008.950659

28. Marowsky A, Burgener J, Falck JR, Fritschy JM, Arand M. Distribution of soluble and microsomal epoxide hydrolase in the mouse brain and its contribution to cerebral epoxyeicosatrienoic acid metabolism. Neuroscience. 2009;163:646–661. doi: 10.1016/j.neuroscience.2009.06.033

29. Wei Q, Doris PA, Pollizotto MV, Boerwinkle E, Jacobs DR, Jr., Siscovick DS, Fornage M. Sequence variation in the soluble epoxide hydrolase gene and subclinical coronary atherosclerosis: interaction with cigarette smoking. Atherosclerosis. 2007;190:26–34. doi: 10.1016/j.atherosclerosis.2006.02.021

30. Zhang W, Koerner IP, Noppens R, Grafe M, Tsai HJ, Morisseau C, Luria A, Hammock BD, Falck JR, Alkayed NJ. Soluble epoxide hydrolase: a novel therapeutic target in stroke. J Cereb Blood Flow Metab. 2007;27:1931–1940. doi: 10.1038/sj.jcbfm.9600494

31. Vacková Š, Kopkan L, Kikerlová S, Husková Z, Sadowski J, Kompanowska-Jezierska E, Hammock BD, Imig JD, Táborský M, Melenovský V, et al. Pharmacological Blockade of Soluble Epoxide Hydrolase Attenuates the Progression of Congestive Heart Failure Combined With Chronic Kidney Disease: Insights From Studies With Fawn-Hooded Hypertensive Rats. Frontiers in Pharmacology. 2019;10. doi: 10.3389/fphar.2019.00018

32. Jiang X-s, Xiang X-y, Chen X-m, He J-l, Liu T, Gan H, Du X-g. Inhibition of soluble epoxide hydrolase attenuates renal tubular mitochondrial dysfunction and ER stress by restoring autophagic flux in diabetic nephropathy. Cell Death & Disease. 2020;11:385. doi: 10.1038/s41419-020-2594-x

33. Ren Q, Ma M, Yang J, Nonaka R, Yamaguchi A, Ishikawa KI, Kobayashi K, Murayama S, Hwang SH, Saiki S, et al. Soluble epoxide hydrolase plays a key role in the pathogenesis of Parkinson’s disease. Proc Natl Acad Sci U S A. 2018;115:E5815–e5823. doi: 10.1073/pnas.1802179115

34. Huang HJ, Wang YT, Lin HC, Lee YH, Lin AM. Soluble Epoxide Hydrolase Inhibition Attenuates MPTP-Induced Neurotoxicity in the Nigrostriatal Dopaminergic System: Involvement of α-Synuclein Aggregation and ER Stress. Molecular neurobiology. 2018;55:138–144. doi: 10.1007/s12035-017-0726-9

35. Rose TE, Morisseau C, Liu JY, Inceoglu B, Jones PD, Sanborn JR, Hammock BD. 1-Aryl-3-(1-acylpiperidin-4-yl)urea inhibitors of human and murine soluble epoxide hydrolase: structure-activity relationships, pharmacokinetics, and reduction of inflammatory pain. J Med Chem. 2010;53:7067–7075. doi: 10.1021/jm100691c

36. Wagner KM, McReynolds CB, Schmidt WK, Hammock BD. Soluble epoxide hydrolase as a therapeutic target for pain, inflammatory and neurodegenerative diseases. Pharmacol Ther. 2017;180:62–76. doi: 10.1016/j.pharmthera.2017.06.006

37. Hammock BD, McReynolds CB, Wagner K, Buckpitt A, Cortes-Puch I, Croston G, Lee KSS, Yang J, Schmidt WK, Hwang SH. Movement to the Clinic of Soluble Epoxide Hydrolase Inhibitor EC5026 as an Analgesic for Neuropathic Pain and for Use as a Nonaddictive Opioid Alternative. Journal of Medicinal Chemistry. 2021;64:1856–1872. doi: 10.1021/acs.jmedchem.0c01886

38. Matin N, Fisher C, Lansdell TA, Hammock BD, Yang J, Jackson WF, Dorrance AM. Soluble epoxide hydrolase inhibition improves cognitive function and parenchymal artery dilation in a hypertensive model of chronic cerebral hypoperfusion. Microcirculation. 2021;28:e12653. doi: 10.1111/micc.12653

39. Hao J, Chen Y, Yao E, Liu X. Soluble epoxide hydrolase inhibition alleviated cognitive impairments via NRG1/ErbB4 signaling after chronic cerebral hypoperfusion induced by bilateral carotid artery stenosis in mice. Brain Res. 2018;1699:89–99. doi: 10.1016/j.brainres.2018.07.002

40. Ghosh A, Comerota MM, Wan D, Chen F, Propson NE, Hwang SH, Hammock BD, Zheng H. An epoxide hydrolase inhibitor reduces neuroinflammation in a mouse model of Alzheimer’s disease. Sci Transl Med. 2020;12. doi: 10.1126/scitranslmed.abb1206

41. Ostermann AI, Herbers J, Willenberg I, Chen R, Hwang SH, Greite R, Morisseau C, Gueler F, Hammock BD, Schebb NH. Oral treatment of rodents with soluble epoxide hydrolase inhibitor 1-(1-propanoylpiperidin-4-yl)-3-[4-(trifluoromethoxy)phenyl]urea (TPPU): Resulting drug levels and modulation of oxylipin pattern. Prostaglandins Other Lipid Mediat. 2015;121:131–137. doi: 10.1016/j.prostaglandins.2015.06.005

42. Islam O, Patil P, Goswami SK, Razdan R, Inamdar MN, Rizwan M, Mathew J, Inceoglu B, Stephen Lee KS, Hwang SH, et al. Inhibitors of soluble epoxide hydrolase minimize ischemia-reperfusion-induced cardiac damage in normal, hypertensive, and diabetic rats. Cardiovasc Ther. 2017;35. doi: 10.1111/1755-5922.12259

43. Wang S, Jiao F, Border JJ, Fang X, Crumpler RF, Liu Y, Zhang H, Jefferson J, Guo Y, Elliott PS, et al. Luseogliflozin, a sodium-glucose cotransporter-2 inhibitor, reverses cerebrovascular dysfunction and cognitive impairments in 18-mo-old diabetic animals. Am J Physiol Heart Circ Physiol. 2022;322:H246–H259. doi: 10.1152/ajpheart.00438.2021

44. Zhang H, Zhang C, Liu Y, Gao W, Wang S, Fang X, Guo Y, Li M, Liu R, Roman RJ, et al. Influence of dual-specificity protein phosphatase 5 on mechanical properties of rat cerebral and renal arterioles. Physiol Rep. 2020;8:e14345. doi: 10.14814/phy2.14345

45. Tang C, Zhang H, Border JJ, Hammock BD, Roman RJ, Fan F. Abstract 15835: Inhibition of Soluble Epoxide Hydrolase Reduces Cognitive Impairment and Improves Cerebral Hemodynamics in AD/ADRD. Circulation. 2023;148:A15835–A15835. doi: doi:10.1161/circ.148.suppl_1.15835

46. Martin M. Cutadapt removes adapter sequences from high-throughput sequencing reads. 2011. 2011;17:3. doi: 10.14806/ej.17.1.200

47. Dobin A, Davis CA, Schlesinger F, Drenkow J, Zaleski C, Jha S, Batut P, Chaisson M, Gingeras TR. STAR: ultrafast universal RNA-seq aligner. Bioinformatics. 2013;29:15–21. doi: 10.1093/bioinformatics/bts635

48. Love MI, Huber W, Anders S. Moderated estimation of fold change and dispersion for RNA-seq data with DESeq2. Genome Biol. 2014;15:550. doi: 10.1186/s13059-014-0550-8

49. Wu T, Hu E, Xu S, Chen M, Guo P, Dai Z, Feng T, Zhou L, Tang W, Zhan L, et al. clusterProfiler 4.0: A universal enrichment tool for interpreting omics data. Innovation (Camb). 2021;2:100141. doi: 10.1016/j.xinn.2021.100141

50. Fan F, Pabbidi MR, Ge Y, Li L, Wang S, Mims PN, Roman RJ. Knockdown of Add3 impairs the myogenic response of renal afferent arterioles and middle cerebral arteries. Am J Physiol Renal Physiol. 2017;312:F971–F981. doi: 10.1152/ajprenal.00529.2016

51. Tang C, Zhang H, Border JJ, Liu Y, Fang X, Jefferson JR, Gregory A, Johnson C, Lee TJ, Bai S, et al. Impact of knockout of dual-specificity protein phosphatase 5 on structural and mechanical properties of rat middle cerebral arteries: implications for vascular aging. Geroscience. 2024;46:3135–3147. doi: 10.1007/s11357-024-01061-y

52. Fan F, Sun CW, Maier KG, Williams JM, Pabbidi MR, Didion SP, Falck JR, Zhuo J, Roman RJ. 20-Hydroxyeicosatetraenoic acid contributes to the inhibition of K+ channel activity and vasoconstrictor response to angiotensin II in rat renal microvessels. PLoS One. 2013;8:e82482. doi: 10.1371/journal.pone.0082482

53. Ge Y, Murphy SR, Fan F, Williams JM, Falck JR, Liu R, Roman RJ. Role of 20-HETE in the impaired myogenic and TGF responses of the Af-Art of Dahl salt-sensitive rats. Am J Physiol Renal Physiol. 2014;307:F509–515. doi: 10.1152/ajprenal.00273.2014

54. Burke M, Pabbidi M, Fan F, Ge Y, Liu R, Williams JM, Sarkis A, Lazar J, Jacob HJ, Roman RJ. Genetic basis of the impaired renal myogenic response in FHH rats. Am J Physiol Renal Physiol. 2013;304:F565–577. doi: 10.1152/ajprenal.00404.2012

55. Fan F, Geurts AM, Murphy SR, Pabbidi MR, Jacob HJ, Roman RJ. Impaired myogenic response and autoregulation of cerebral blood flow is rescued in CYP4A1 transgenic Dahl salt-sensitive rat. Am J Physiol Regul Integr Comp Physiol. 2015;308:R379–390. doi: 10.1152/ajpregu.00256.2014

56. Faraci FM, Heistad DD. Regulation of large cerebral arteries and cerebral microvascular pressure. Circ Res. 1990;66:8–17. doi: 10.1161/01.res.66.1.8

57. Mayhan WG, Heistad DD. Role of veins and cerebral venous pressure in disruption of the blood-brain barrier. Circ Res. 1986;59:216–220. doi: 10.1161/01.res.59.2.216

58. Pires PW, Dabertrand F, Earley S. Isolation and Cannulation of Cerebral Parenchymal Arterioles. J Vis Exp. 2016. doi: 10.3791/53835

59. Claassen J, Thijssen DHJ, Panerai RB, Faraci FM. Regulation of Cerebral Blood Flow in Humans: Physiology and Clinical Implications of Autoregulation. Physiol Rev. 2021. doi: 10.1152/physrev.00022.2020

60. Shekhar S, Wang S, Mims PN, Gonzalez-Fernandez E, Zhang C, He X, Liu CY, Lv W, Wang Y, Huang J, et al. Impaired Cerebral Autoregulation-A Common Neurovascular Pathway in Diabetes may Play a Critical Role in Diabetes-Related Alzheimer’s Disease. Curr Res Diabetes Obes J. 2017;2:555587.

61. Administration USFaD. FDA approves treatment for adults with Alzheimer’s disease. U.S. Food and Drug Administration. https://www.fda.gov/drugs/news-events-human-drugs/fda-approves-treatment-adults-alzheimers-disease. 2024.

62. Withington CG, Turner RS. Amyloid-Related Imaging Abnormalities With Anti-amyloid Antibodies for the Treatment of Dementia Due to Alzheimer’s Disease. Front Neurol. 2022;13:862369. doi: 10.3389/fneur.2022.862369

63. Gaugler J, James B, Johnson T, Marin A, Weuve J, Assoc As. 2019 Alzheimer’s disease facts and figures. Alzheimers Dement. 2019;15:321–387. doi: 10.1016/j.jalz.2019.01.010

64. Samaras K, Lutgers HL, Kochan NA, Crawford JD, Campbell LV, Wen W, Slavin MJ, Baune BT, Lipnicki DM, Brodaty H, et al. The impact of glucose disorders on cognition and brain volumes in the elderly: the Sydney Memory and Ageing Study. Age (Dordr). 2014;36:977–993. doi: 10.1007/s11357-013-9613-0

65. Wang S, Mims PN, Roman RJ, Fan F. Is Beta-Amyloid Accumulation a Cause or Consequence of Alzheimer’s Disease? J Alzheimers Parkinsonism Dement. 2016;1.

66. Imig JD. Epoxyeicosatrienoic Acids and 20-Hydroxyeicosatetraenoic Acid on Endothelial and Vascular Function. Adv Pharmacol. 2016;77:105–141. doi: 10.1016/bs.apha.2016.04.003

67. Moghaddam MF, Grant DF, Cheek JM, Greene JF, Williamson KC, Hammock BD. Bioactivation of leukotoxins to their toxic diols by epoxide hydrolase. Nat Med. 1997;3:562–566. doi: 10.1038/nm0597-562

68. Hildreth K, Kodani SD, Hammock BD, Zhao L. Cytochrome P450-derived linoleic acid metabolites EpOMEs and DiHOMEs: a review of recent studies. J Nutr Biochem. 2020;86:108484. doi: 10.1016/j.jnutbio.2020.108484

69. Griñán-Ferré C, Codony S, Pujol E, Yang J, Leiva R, Escolano C, Puigoriol-Illamola D, Companys-Alemany J, Corpas R, Sanfeliu C, et al. Pharmacological Inhibition of Soluble Epoxide Hydrolase as a New Therapy for Alzheimer’s Disease. Neurotherapeutics. 2020;17:1825–1835. doi: 10.1007/s13311-02000854-1

70. Lee HT, Lee KI, Chen CH, Lee TS. Genetic deletion of soluble epoxide hydrolase delays the progression of Alzheimer’s disease. J Neuroinflammation. 2019;16:267. doi: 10.1186/s12974-019-1635-9

71. Sun CP, Zhang XY, Zhou JJ, Huo XK, Yu ZL, Morisseau C, Hammock BD, Ma XC. Inhibition of sEH via stabilizing the level of EETs alleviated Alzheimer’s disease through GSK3β signaling pathway. Food Chem Toxicol. 2021;156:112516. doi: 10.1016/j.fct.2021.112516

72. Yu D, Hennebelle M, Sahlas DJ, Ramirez J, Gao F, Masellis M, Cogo-Moreira H, Swartz RH, Herrmann N, Chan PC, et al. Soluble Epoxide Hydrolase-Derived Linoleic Acid Oxylipins in Serum Are Associated with Periventricular White Matter Hyperintensities and Vascular Cognitive Impairment. Transl Stroke Res. 2019;10:522–533. doi: 10.1007/s12975-018-0672-5

73. Fan F, Simino J, Auchus AP, Knopman DS, Boerwinkle E, Fornage M, Mosley TH, Roman RJ. Functional variants in CYP4A11 and CYP4F2 are associated with cognitive impairment and related dementia endophenotypes in the elderly. In: The 16th International Winter Eicosanoid Conference. Baltimore; 2016:CV5.

74. Liang Z, Zhang B, Xu M, Morisseau C, Hwang SH, Hammock BD, Li QX. 1-Trifluoromethoxyphenyl-3-(1-propionylpiperidin-4-yl) Urea, a Selective and Potent Dual Inhibitor of Soluble Epoxide Hydrolase and p38 Kinase Intervenes in Alzheimer’s Signaling in Human Nerve Cells. ACS Chem Neurosci. 2019;10:4018–4030. doi: 10.1021/acschemneuro.9b00271

75. Minaz N, Razdan R, Hammock BD, Goswami SK. An inhibitor of soluble epoxide hydrolase ameliorates diabetes-induced learning and memory impairment in rats. Prostaglandins Other Lipid Mediat. 2018;136:84–89. doi: 10.1016/j.prostaglandins.2018.05.004

76. Pardeshi R, Bolshette N, Gadhave K, Arfeen M, Ahmed S, Jamwal R, Hammock BD, Lahkar M, Goswami SK. Docosahexaenoic Acid Increases the Potency of Soluble Epoxide Hydrolase Inhibitor in Alleviating Streptozotocin-Induced Alzheimer’s Disease-Like Complications of Diabetes. Front Pharmacol. 2019;10:288. doi: 10.3389/fphar.2019.00288

77. Wu J, Fan Z, Zhao Y, Chen Q, Xiao Q. Inhibition of soluble epoxide hydrolase (sEH) protects hippocampal neurons and reduces cognitive decline in type 2 diabetic mice. Eur J Neurosci. 2021;53:2532–2540. doi: 10.1111/ejn.15150

78. Luo J, Hu S, Fu M, Luo L, Li Y, Li W, Cai Y, Dong R, Yang Y, Tu L, et al. Inhibition of soluble epoxide hydrolase alleviates insulin resistance and hypertension via downregulation of SGLT2 in the mouse kidney. J Biol Chem. 2021;296:100667. doi: 10.1016/j.jbc.2021.100667

79. Luria A, Bettaieb A, Xi Y, Shieh GJ, Liu HC, Inoue H, Tsai HJ, Imig JD, Haj FG, Hammock BD. Soluble epoxide hydrolase deficiency alters pancreatic islet size and improves glucose homeostasis in a model of insulin resistance. Proc Natl Acad Sci U S A. 2011;108:9038–9043. doi: 10.1073/pnas.1103482108

80. Zhang LN, Vincelette J, Chen D, Gless RD, Anandan SK, Rubanyi GM, Webb HK, MacIntyre DE, Wang YX. Inhibition of soluble epoxide hydrolase attenuates endothelial dysfunction in animal models of diabetes, obesity and hypertension. Eur J Pharmacol. 2011;654:68–74. doi: 10.1016/j.ejphar.2010.12.016

81. Luther JM, Ray J, Wei D, Koethe JR, Hannah L, DeMatteo A, Manning R, Terker AS, Peng D, Nian H, et al. GSK2256294 Decreases sEH (Soluble Epoxide Hydrolase) Activity in Plasma, Muscle, and Adipose and Reduces F2-Isoprostanes but Does Not Alter Insulin Sensitivity in Humans. Hypertension. 2021;78:1092–1102. doi: 10.1161/hypertensionaha.121.17659

82. Liu R, Hossain MM, Chen X, Jin JP. Mechanoregulation of SM22α/Transgelin. Biochemistry. 2017;56:5526–5538. doi: 10.1021/acs.biochem.7b00794

83. Shao H, Wang JH, Pollak MR, Wells A. α-actinin-4 is essential for maintaining the spreading, motility and contractility of fibroblasts. PLoS One. 2010;5:e13921. doi: 10.1371/journal.pone.0013921

84. Gao W, Liu Y, Fan L, Zheng B, Jefferson JR, Wang S, Zhang H, Fang X, Nguyen BV, Zhu T, et al. Role of γ-adducin in actin cytoskeleton rearrangements in podocyte pathophysiology. Am J Physiol Renal Physiol. 2021;320:F97–f113. doi: 10.1152/ajprenal.00423.2020

85. Lader AS, Kwiatkowski DJ, Cantiello HF. Role of gelsolin in the actin filament regulation of cardiac L-type calcium channels. Am J Physiol. 1999;277:C1277–1283. doi: 10.1152/ajpcell.1999.277.6.C1277

86. Yu Q, Wu C, Chen Y, Li B, Wang R, Huang R, Li X, Gu D, Wang X, Duan X, et al. Inhibition of LIM kinase reduces contraction and proliferation in bladder smooth muscle. Acta Pharm Sin B. 2021;11:1914–1930. doi: 10.1016/j.apsb.2021.01.005

87. Sun J, Qiao YN, Tao T, Zhao W, Wei LS, Li YQ, Wang W, Wang Y, Zhou YW, Zheng YY, et al. Distinct Roles of Smooth Muscle and Non-muscle Myosin Light Chain-Mediated Smooth Muscle Contraction. Front Physiol. 2020;11:593966. doi: 10.3389/fphys.2020.593966

88. Fisher SA. Vascular smooth muscle phenotypic diversity and function. Physiol Genomics. 2010;42a:169–187. doi: 10.1152/physiolgenomics.00111.2010

89. Lin Z, Shi JL, Chen M, Zheng ZM, Li MQ, Shao J. CCL2: An important cytokine in normal and pathological pregnancies: A review. Front Immunol. 2022;13:1053457. doi: 10.3389/fimmu.2022.1053457

90. Munguia ME, Govezensky T, Martinez R, Manoutcharian K, Gevorkian G. Identification of amyloid-beta 1-42 binding protein fragments by screening of a human brain cDNA library. Neurosci Lett. 2006;397:79–82. doi: 10.1016/j.neulet.2005.11.061

91. Reza K, Mostafa R-T, Bahareh T, Sadegh A. Multi-regional Neurodegeneration in Alzheimer’s disease: Meta-analysis and data integration of transcriptomics data. bioRxiv. 2018:245571. doi: 10.1101/245571

92. Adav SS, Park JE, Sze SK. Quantitative profiling brain proteomes revealed mitochondrial dysfunction in Alzheimer’s disease. Mol Brain. 2019;12:8. doi: 10.1186/s13041-019-0430-y

93. Bianchi M, Crinelli R, Arbore V, Magnani M. Induction of ubiquitin C (UBC) gene transcription is mediated by HSF1: role of proteotoxic and oxidative stress. FEBS Open Bio. 2018;8:1471–1485. doi: 10.1002/2211-5463.12484

94. Cohen RM, Rezai-Zadeh K, Weitz TM, Rentsendorj A, Gate D, Spivak I, Bholat Y, Vasilevko V, Glabe CG, Breunig JJ, et al. A transgenic Alzheimer rat with plaques, tau pathology, behavioral impairment, oligomeric aβ, and frank neuronal loss. J Neurosci. 2013;33:6245–6256. doi: 10.1523/jneurosci.3672-12.2013

95. Chiamvimonvat N, Ho CM, Tsai HJ, Hammock BD. The soluble epoxide hydrolase as a pharmaceutical target for hypertension. Journal of cardiovascular pharmacology. 2007;50:225–237. doi: 10.1097/FJC.0b013e3181506445

96. Cipolla MJ, Porter JM, Osol G. High Glucose Concentrations Dilate Cerebral Arteries and Diminish Myogenic Tone Through an Endothelial Mechanism. Stroke. 1997;28:405–411. doi: doi:10.1161/01.STR.28.2.405

97. O’Brien CE, Santos PT, Kulikowicz E, Lee JK, Koehler RC, Martin LJ. Neurologic effects of short-term treatment with a soluble epoxide hydrolase inhibitor after cardiac arrest in pediatric swine. BMC Neuroscience. 2020;21:43. doi: 10.1186/s12868-020-00596-y

